# Efficient Integrative Multi-SNP Association Analysis using Deterministic Approximation of Posteriors

**DOI:** 10.1101/026450

**Authors:** Xiaoquan Wen, Yeji Lee, Francesca Luca, Roger Pique-Regi

## Abstract

With the increasing availability of functional genomic data,^1–3^ incorporating genomic annotations into genetic association analysis has become a standard procedure. However, the existing methods often lack rigor and/or computational efficiency and consequently do not maximize the utility of functional annotations. In this paper, we propose a rigorous inference procedure to perform integrative association analysis incorporating genomic annotations for both traditional GWAS and emerging molecular QTL mapping studies. In particular, we propose an algorithm, named “Deterministic Approximation of Posteriors” (DAP), which enables highly efficient and accurate joint enrichment analysis and identification of multiple causal variants. We use a series of simulation studies to highlight the power and computational efficiency of our proposed approach and further demonstrate it by analyzing the cross-population eQTL data from the GEUVADIS project and the multi-tissue eQTL data from the GTEx project. In particular, we find that genetic variants predicted to disrupt transcription factor binding sites are enriched in *cis*-eQTLs across all tissues. Moreover, the enrichment estimates obtained across the tissues are correlated with the cell types for which the annotations are derived.

## Introduction

Association analysis has become a powerful tool for identifying genetic variants that impact complex traits at both the organismal and molecular levels: in the past decade, genome-wide association studies (GWAS) have successfully identified a rich catalog of genetic variants that are linked to many human diseases. Most recently, molecular QTL mapping has revealed an abundance of quantitative trait loci (QTLs) for cellular phenotypes such as gene expression,^3,4^ chromatin accessibility,^5^ histone modifications^6^ and DNA methylation.^7^ Nevertheless, the causal molecular pathways from genetic variants to complex phenotypes remain poorly understood.^8^ This is mainly because a good proportion of identified trait-associated variants are located in the non-coding regions of the genome, and our knowledge of the functional roles of non-coding variants is generally lacking. With the recent advancements in high-throughput experimental technologies, functional annotations for regulatory variants have become increasingly available.^1–3^ As a consequence, it is now feasible to perform association analysis incorporating functional genomic annotations. The integrative analysis strategy presents two obvious advantages: first, it improves the power of association analysis by prioritizing functional variants; second, it helps to reveal the underlying molecular mechanisms that lead to the observed associations.

In the past, integrative analysis was typically performed by searching for overlaps between putative association signals and SNP annotations. This analysis strategy implicitly assumes that a SNP with specific genomic annotations is likely causal. To justify the results from the posthoc overlapping analysis, quantitatively validating this implicit assumption from the observed association data, which essentially requires estimating the enrichment levels of the annotations in the association signals, is critical. This point becomes particularly crucial when multiple types of annotations are used, and a rigorous quantitative enrichment analysis should help to determine which annotations are relevant and how much we should weigh each annotation. The availability of functional annotations also enables high-resolution multi-SNP genetic association analysis. From both GWAS and molecular QTL mapping studies, it is increasingly evident that multiple independent association signals can co-exist in a relatively small genomic region. Multi-SNP fine-mapping analysis has now become a standard procedure to tease out potential multiple association signals. It is only natural that genomic annotations are integrated into this process.

Recently, a few computational approaches for integrative enrichment and association analysis have been proposed and successfully demonstrated in molecular QTL mapping,^10^ and GWAS.^11,12^ However, these existing approaches make simplifying assumptions for either enrichment analysis^12^ or multi-SNP fine-mapping analysis.^9,11^ Therefore, the power of integrative analysis has not been maximized and can be further improved. In addition, computational efficiency has always been a hurdle in terms of applying a probabilistic integrative analysis approaches to genetic data at the genome-wide scale.

In this paper, we propose a probabilistic hierarchical model that is generalized from our recent work^13^ to describe multi-SNP genetic associations while accounting for functional genomic annotations. Based on this model, we consider analyzing genetic association data in two settings: traditional GWAS and molecular *cis*-QTL mapping studies. Note that a distinct feature of molecular QTL mapping is that tens of thousands (or hundreds of thousands) of molecular phenotypes (e.g., gene expression, DNA methylation) are simultaneously measured and analyzed, which imposes some unique statistical challenges. In addition, the candidate genomic region for each molecular phenotype is typically defined in the proximity of relevant genomic landmarks of the corresponding molecular phenotypes (e.g., transcription start site of a target gene for expression phenotypes) and much smaller in length (usually spanning 1 to 2 Mb) comparing to GWAS. We outline a 3-stage inference procedure to sequentially perform enrichment analysis, QTL discovery and multi-SNP fine-mapping. One of our main contributions is a computationally efficient algorithm for Bayesian multi-SNP association analysis. This fast fitting algorithm, named Deterministic Approximation of Posteriors (DAP), facilitates the proposed rigorous integrative inference procedure. Compared to the alternative fitting algorithm, i.e., the Markov Chain Monte Carlo (MCMC) algorithm, we show that the DAP is several hundreds times faster and more accurate for genetic association analysis. Taking full advantage of the DAP algorithm, we lay out the analytic strategies for analyzing genetic association data from GWAS and molecular *cis*-QTL mapping studies, and we demonstrate the proposed procedures through a series of simulation studies and real data applications.

## Methods

### Model and Notation

First, we consider a generic setting of association analysis of a single quantitative trait and *p* SNPs both measured for *n* unrelated individuals. We model the genotype-phenotype association using a multiple linear regression model,

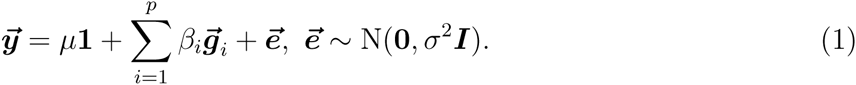

For each SNP *i*, we denote its binary association status, *γ_i_*, by dichotomizing its corresponding genetic effect *β_i_*, i.e, *γ_i_* = 1 if *β_i_* ≠ 0 and 0 otherwise. In particular, we refer to the causal SNPs for which *γ_i_* = 1 as the quantitative trait nucleotides (QTNs).^9^ Our primary interest for association analysis is the inference of 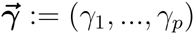. To integrate genomic annotation into the association analysis, we assume that having certain genomic features will increase (or decrease) the odds that a particular SNP is a QTN. Equivalently, certain genomic features are enriched (or depleted) in QTNs. We quantitatively represent this assumption using an *a priori* independent logistic model for each *γ_i_*, i.e.,

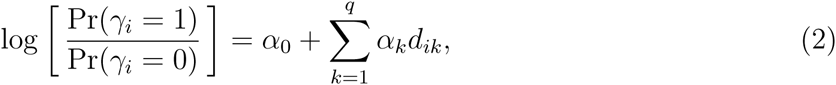

where 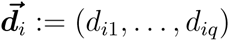 denotes *q* genomic annotations that are specific to SNP *i* at a particular locus and *α*_1_, …, *α_q_* are referred to as the *enrichment parameters*. Note that the annotations can be either categorical or continuous in this framework. We assume that the phenotype data, 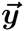, the genotype data, 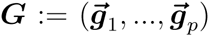, and the annotation data, 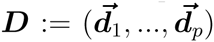, are observed, while the enrichment parameters, 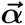, are unknown.

For molecular QTL mapping, tens of thousands of phenotypes are simultaneously measured, and we denote the collection of all measured phenotypes by 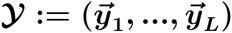. For each phenotype, a small genomic region, typically spanning 1 to 2 Mb and on average containing a few thousands of SNPs, is pre-defined as the candidate locus in the proximity of relevant genomic landmarks of the corresponding molecular phenotypes, and we denote the union of the SNP genotypes from all candidate loci by ***G*** := (*G*_1_,…, *G_L_*). Similarly, we use ***D*** := (*D*_1_,…, *D_L_*) and 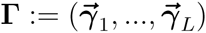 to denote the collections of annotations and latent association status, respectively.

In GWAS, there is usually only one phenotype of interest, which can be viewed as a special case of molecular QTL mapping. Nevertheless, it is important to note that the candidate region for GWAS spans the whole genome.

### Inference Procedure

We propose an inference procedure consisting of three inter-related stages to fit the proposed hierarchical model. Sequentially, these stages are as follows:

1. estimating the enrichment parameter 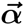 using the full data 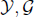 and 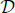 for enrichment analysis
2. screening candidate loci for QTL discovery
3. performing multi-SNP fine-mapping for the high-priority loci identified in stage 2

The maximum likelihood estimate (MLE) of 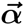 can be obtained by the EM algorithm proposed in our recent work.^13^ Briefly, the EM algorithm treats **Γ** as missing data and pools information across all available loci. In the E-step, the posterior inclusion probability (PIP) for each SNP *i* at each locus *l* (namely, 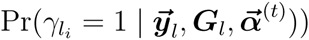 is computed given the current estimate of 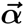; in the M-step, a logistic regression model is fit by plugging in the PIPs as the response variables and SNP annotations as predictors. The estimate of 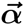 is subsequently updated by the corresponding fitted regression coefficients.

Given the MLE of the enrichment parameter, 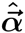, we then attempt to identify genomic loci that are likely to harbor causal QTNs. This is achieved by testing the null hypothesis, 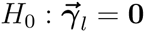, for each candidate locus *l* using a Bayesian false discovery rate (FDR) control procedure. Specifically, the null hypothesis is rejected if the locus-level posterior probability 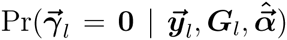 is smaller than the pre-defined threshold determined by the observed data and desired FDR control level.^14^ At the end of this stage, we gather a list of potential QTLs for fine-mapping.

Finally, we perform multi-SNP fine-mapping analysis for the identified QTLs. In particular, we compute the posterior distribution for each locus *l*, namely, 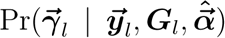, to i) identify potentially multiple independent association signals within locus *l* and ii) assess the importance of each SNP by computing its PIP, i.e., 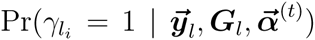. A credible set of potential causal SNPs for each independent signal can then be constructed from the resulting PIPs in a manner similar to previously proposed methods.^13,15^ This Bayesian approach for multi-SNP analysis has been known to present some unique advantages over the traditional conditional analysis approach. For example, it fully accounts for patterns of linkage disequilibrium (LD) and shows superior power in discovering independent association signals.^13,16^

This 3-stage procedure represents a coherent empirical Bayes strategy to fit the proposed hierarchical model for inference. In all three stages, the computational difficulty lies in the efficient evaluation of the posterior probability 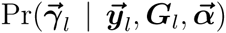. We propose an algorithm to tackle this problem in the following sections. The software package implementing the computational approaches (in C** programming language) is freely available (Web Resources).

### Deterministic Approximation of Posteriors

The computation of the target posterior probability 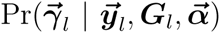 is conceptually straightforward by applying the Bayes theorem, i.e.,

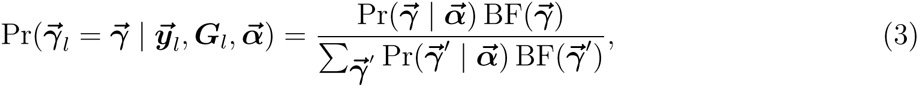

where the Bayes factor

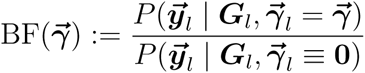

represents the marginal likelihood function of 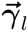 evaluated at 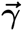. Based on (3), the PIP of each candidate SNP can be subsequently marginalized from 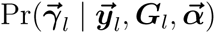.

For any given 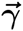 value, both the Bayes factor (whose computation involves integrating out the the nuisance parameters *μ, β* and *σ*^2^) and the prior probability can be analytically evaluated.^17,18^ The difficulty lies in evaluating the normalizing constant

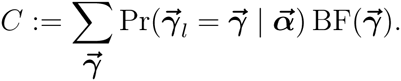

For a locus consisting of p candidate SNPs, the exact computation requires enumerating all 2^*p*^ possible 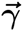 values; hence, it is intractable even for modest *p*. Previously, the only feasible solution was to employ a Markov Chain Monte Carlo (MCMC) algorithm.^13,16,19^ However, the MCMC algorithm is computationally too costly in our grand scheme for integrative genetic association analysis: the evaluation of 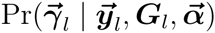 for every locus is required for each E-step in the EM algorithm for enrichment analysis. Furthermore, the inherent stochastic variation in the MCMC algorithm may affect the performance and reproducibility of the overall analysis.

Here, we present an alternative algorithm to perform deterministic approximation of posteriors (DAP) for each locus and efficiently compute PIPs for all candidate SNPs. This algorithm is mainly motivated by two observations in genetic association analysis. First, in almost all genetic applications, the number of convincing QTLs (i.e., those have relatively large effect sizes) discovered from the association data are typically small compared with the number of candidate SNPs within a candidate locus (typically 1 to 2 Mb). In molecular QTL mapping, this observation is also supported by many recent experimental work.^20–22^ It implies that the vast majority of the posterior probability mass in the space of all possible combinations of SNPs must be concentrated in a much lower-dimensional subspace. That is, only association models containing a few SNPs are likely to have non-negligible posterior probabilities within a locus. Second, noteworthy QTL SNPs, as reflected by their non-negligible PIP values, are thought to typically show modest to strong marginal association signals in either single-SNP or conditional analysis. Based on the above observations, we design the DAP algorithm to adaptively select a small subset of noteworthy candidate QTL SNPs and thoroughly explore the low-dimensional model space composed by these SNPs within each candidate locus. In addition, the DAP algorithm applies a combinatorial approximation to estimate the posterior probability mass from the unexplored model space. Unlike the MCMC, the DAP algorithm is highly parallelizable, and our implementation takes full advantage of this property. More specifically, the proposed DAP algorithm approximates the normalizing constant *C* by

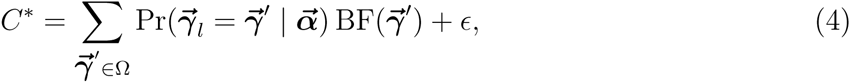

where Ω denotes a subset of the selected most plausible models to be explored explicitly and e is an estimate of the approximation error 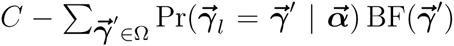. The key to the DAP algorithm is the construction of the set Ω: it is desirable that models in Ω capture the vast majority of the posterior probability mass; on the other hand, Ω should be compact enough for efficient exploration. In this paper, we propose two different approaches to construct Ω. In both cases, we define the size of the association model, 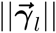, as the number of assumed QTNs (also known as the 0-norm of the vector 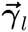), i.e., 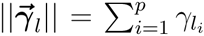, and partition the complete model space of 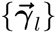 by the size of association models, i.e., 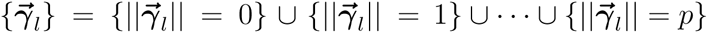.

### Adaptive DAP Algorithm

The first approach, named adaptive DAP, includes the null model and all the single SNP association models in the candidate set Ω. For a larger size of candidate models, it approximates 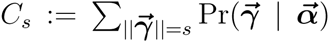 by a corresponding estimate 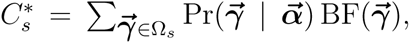, where Ω_*s*_ consists of a subset of association models with size s but is constructed only from a set of adaptively selected high-priority SNPs. The adaptive selection of the high-priority SNPs is similar to a Bayesian version of conditional analysis^23^ that naturally accounts for LD. More specifically, suppose that a “best” model with the maximum posterior probability for 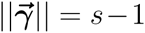 has been identified. The SNP selection procedure then goes through all candidate SNPs, adding a single SNP at a time to the existing best model, and evaluates their posterior probabilities of being the sole additional QTN (see the details in the Appendix A). Note that this procedure is similar to single-SNP analysis and is computationally trivial. The candidate SNPs whose posterior probabilities in the conditional analysis are greater than a pre-defined threshold λ, which is a valid probability measure (by default, we set λ = 0.01), are then added to the existing subset of high-priority SNPs. Finally, the DAP algorithm enumerates the updated subset of priority SNPs for all combinations of 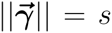 to compute 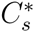 and, in the process, records the “best” posterior model with the increased model size.

Additionally, the adaptive DAP only extensively explores the model partitions with relatively small sizes. Suppose that there are truly *K* QTLs in *p* candidate SNPs. It should be clear that {*C_s_* } becomes a (sharply) decreasing sequence as *s* > *K* and that the behavior of this decreasing sequence is mathematically predictable (Appendix B). This behavior occurs because the marginal likelihood becomes saturated as the model size exceeds the number of true associations and because the additional prior term imposes a hefty penalty on the overall product. Utilizing this fact, we derive an approximate recursive relationship between *C_s_* and *C*_*s*\*1_ as *s* ≥ *K* (Appendix B). Based on this relationship, the stopping rule for explicit exploration is determined, and we estimate *ϵ* by

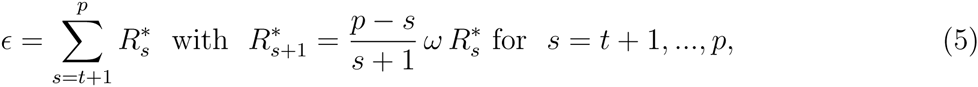

where *t* is the stopping point of the extensive exploration, 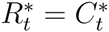, and 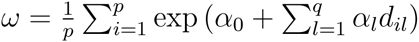 represents the average prior odds ratio across SNPs. This estimation essentially assumes that the marginal likelihood is completely saturated for the partitions with *s* > *t*, and the overall contribution to the normalizing constant from each size partition can be roughly estimated by re-calibrating the prior changes (see the details in Appendix B). To ensure a high accuracy for the approximation, we also build in an *optional* criterion on top of the stopping rule by monitoring the convergence of the partial sum 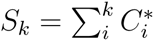 and enforcing the exploration until

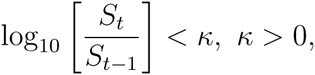

or, equivalently 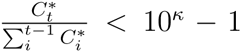. By default, we set *κ* = 0.01. This additional criterion only makes a difference for the partitions whose model sizes barely exceed the estimated size of the saturated models: instead of using the combinatorial estimate of the corresponding 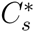, it enforces additional DAP explorations for more accurate evaluations.

Finally, it should be recognized that the built-in tuning parameters (*λ, κ*) enable great flexibility to run the adaptive DAP. As both *λ* → 0 and *κ* → 0, the adaptive DAP enumerates *all* models and becomes an *exact* calculation with no loss of precision, whereas when *λ* is very large, the behavior of the DAP algorithm becomes very similar to the commonly applied step-wise conditional analysis that has very high computational efficiency. In practice, we attempt to strike a good balance between the precision and efficiency.

### DAP-K Algorithm

Instead of adaptively selecting a subset of high-priority SNPs from all the model size partitions, the DAP algorithm can also be applied by pre-fixing the maximum model size (namely, *K*) while allowing the exploration of all possible SNP combinations under the restriction. We refer to this variant of the algorithm as the DAP-*K* algorithm. In the special case of *K* = 1 (DAP-1), the algorithm essentially assumes that at most one causal QTL exists in the region of interest. Although this very assumption has been successfully utilized by many other approaches,^9,11,17,23^ it has always been formulated as an explicit prior assumption and hence requires a somewhat non-natural parameterization that also complicates the maximization step when used in the EM algorithm for enrichment analysis (Appendix C). The DAP-1 algorithm provides the advantage of considerably faster computation, even when compared with the adaptive version of the DAP algorithm. More importantly, it can be applied using only summary statistics from single-SNP association analysis (in the form of the marginal estimate of the genetic effect and its standard error for each SNP). This feature is particularly attractive, especially when the individual-level genotype and phenotype information is difficult to access. We provide the derivation and other technical details for the DAP-*K* algorithm in the Appendix C.

### Applying DAP in Inference

We use both variants of the DAP algorithms in our inference procedure. Specifically, we propose applying the DAP-1 algorithm in the EM algorithm for enrichment analysis and the adaptive DAP for multi-SNP fine-mapping at the last stage.

The performance of the enrichment analysis mostly relies on the *average* accuracy of the PIP estimates. We show, both theoretically (Appendix E) and numerically (Figure 4), that the DAP-1 algorithm provides on average precise estimates suitable for enrichment analysis. Most importantly, the DAP-1 algorithm exhibits the best computational efficiency among the appropriate alternatives (e.g., adaptive DAP, MCMC).

For the multi-SNP analysis in the final fine-mapping stage, we strongly recommend applying the adaptive DAP algorithm. Although the DAP-1 algorithm only yields inferior results for a small proportion of the loci that harbor multiple QTNs, we argue that identifying multiple independent association signals from those loci is of particular importance for the overall analysis. To achieve better accuracy for all loci, the adaptive DAP seems a logical choice for multi-SNP fine-mapping analysis.

### Application to GWAS

In practice, the DAP works well for small genomic regions harboring a handful of QTNs. This is typically the case in molecular QTL mapping, where candidate loci usually span no more than 2 Mb. When there are more QTNs (e.g., > 5) in a locus, the adaptive DAP exploration with high precision may become time consuming because the size of the candidate set Ω grows exponentially fast with the increasing number of independent signals. Nevertheless, in applications of GWAS, we essentially consider a single locus that spans the whole genome, and for a single trait, the number of independent association signals may range from hundreds to thousands.

To apply the DAP to GWAS (or molecular QTL mapping with considerably larger candidate loci), we propose an additional approximation that factorizes 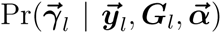 (where locus *l* spans a much larger genomic region) into

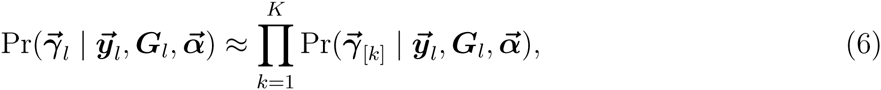

where 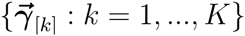 represents a partition of 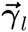 by sets of non-overlapping LD blocks. This factorization is based on previous theoretical results.^18,24^ Recently, Berisa and Pickrell^25^ provided a working recipe to segment the full genome based on the population-specific LD structures. Based on these results, we provide mathematical arguments to justify the factorization (Appendix D). Briefly, applying the analytic approximation of the Bayes factors,^18^ it can be shown that

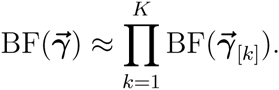

This result, along with the fact that our priors are independent across SNPs, naturally leads to the approximate factorization of the posterior probability. As an important consequence, the factorization (6) suggests that the DAP can be applied to each LD block independently.

## Results

First, we perform a series of simulation studies to examine the accuracy and efficiency of the proposed DAP algorithms in our inference procedure. We then apply the proposed approach to analyze two large-scale eQTL data sets.

### Simulation Studies

#### Enrichment Analysis with DAP

The integration of DAP into the EM algorithm enables the efficient estimation of enrichment parameters using large-scale QTL data sets. To investigate the performance of the enrichment analysis, we simulate a modest-scale eQTL data set to mimic the genome-wide investigation of *cis*-eQTLs. Specifically in each simulation, we select a subset of 1,500 random genes from the GEUVADIS data.^4^ For each gene, the real genotypes of 50 *cis*-SNPs from 343 European individuals are used in the simulation. We annotate 20% of the SNPs with a binary feature. For each SNP, we determine its binary association status by performing a Bernoulli trial with the success rate 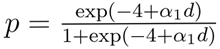. Given the QTNs, we then simulate the expression levels according to a multiple linear regression model with residual error variance set to 1. More specifically, the genetic effect of each QTN is drawn from an independent normal distribution N(0, 0.6^2^). As a result, the simulated data sets resemble the practically observed *cis*-eQTL data (Figure 1). We vary the *α*_1_ values from 0.00 to 1.00, and we generate 100 data sets for each *α*_1_ value.

**Figure 1:**
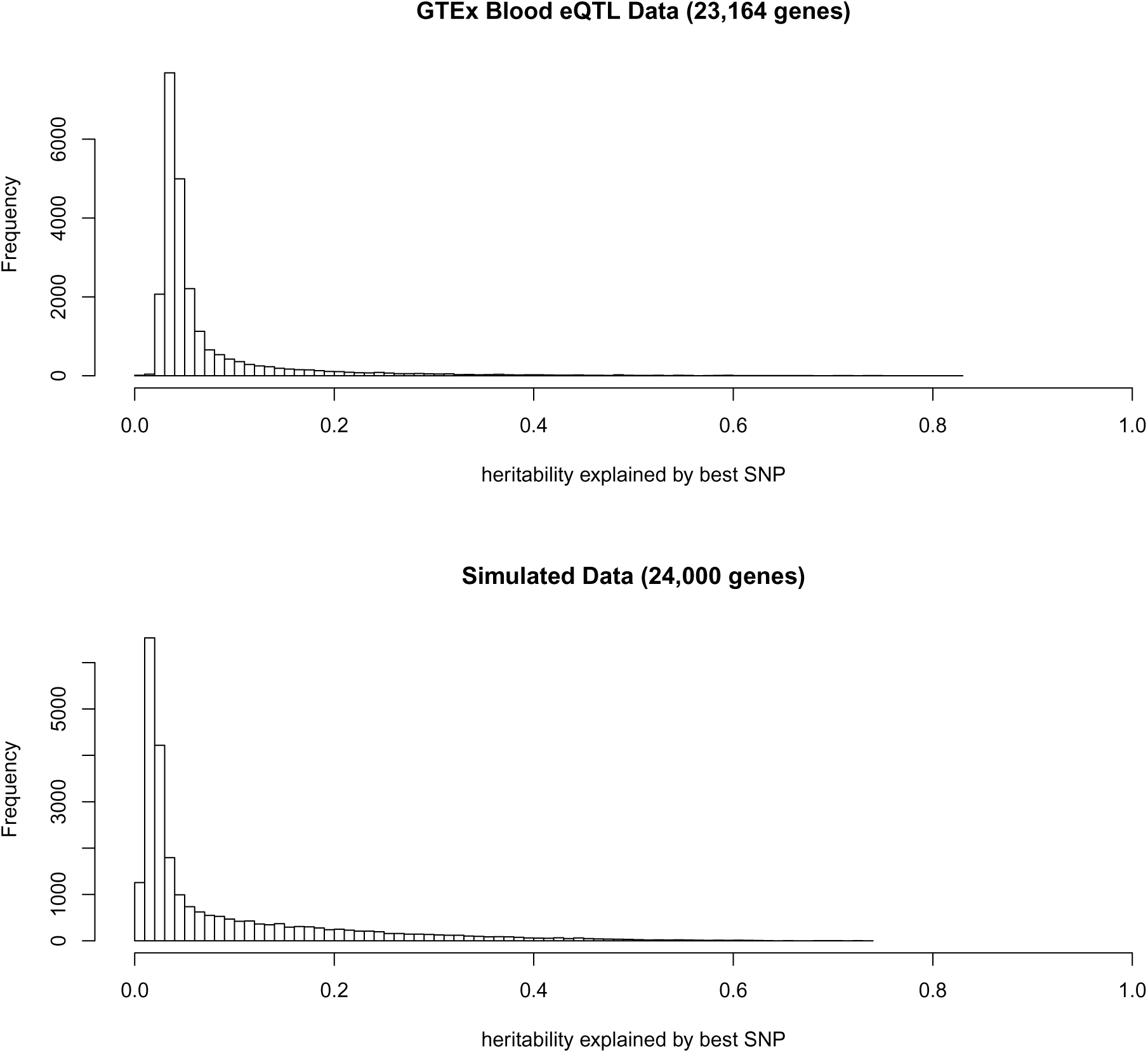
Comparison of simulated data set with the actual GTEx whole blood *cis*-eQTL data.

For each gene in each data set, we find the best associated SNP based on single-SNP testing, and compute the heritability explained by the best SNP using a simple linear regression model. The histograms show the distribution of the heritability across all genes. The similarity of the two histograms indicates that the simulated data sets closely resemble the real observed *cis*-eQTL data.

We analyze the simulated data sets using two different implementations of the EM algorithm with the E-step approximated by the DAP-1 and the adaptive DAP. For evaluation, we also estimate *α*_1_ by fitting a logistic regression model using the true association status of each SNP. This analysis represents a theoretical best-case scenario, and its results should be regarded as the bound of the most optimal outcome from any analysis that infers the latent association status (Γ) from observed data.

Figure 2 shows that the estimates from the adaptive DAP and DAP-1 are both seemingly unbiased. As expected, the variability of the point estimates from both DAP implementations is higher than that from the best-case method because of the uncertainty in determining the true association status of each SNP. The estimates of the 95% confidence intervals from the individual simulations also confirm this finding (Figure 3). Although the adaptive DAP seemingly generates more accurate estimates on average, we conclude that the numerical performance of DAP-1 is very comparable. Importantly, DAP-1 provides superior computational efficiency: the average running time for the DAP-1-embedded EM algorithm (with 10 parallel threads in the E-step) is 65.05 seconds; in comparison, the adaptive DAP-embedded EM runs for 387.30 seconds on average (which is a combination of slightly longer iterations and longer running times per iteration).

**Figure 2:**
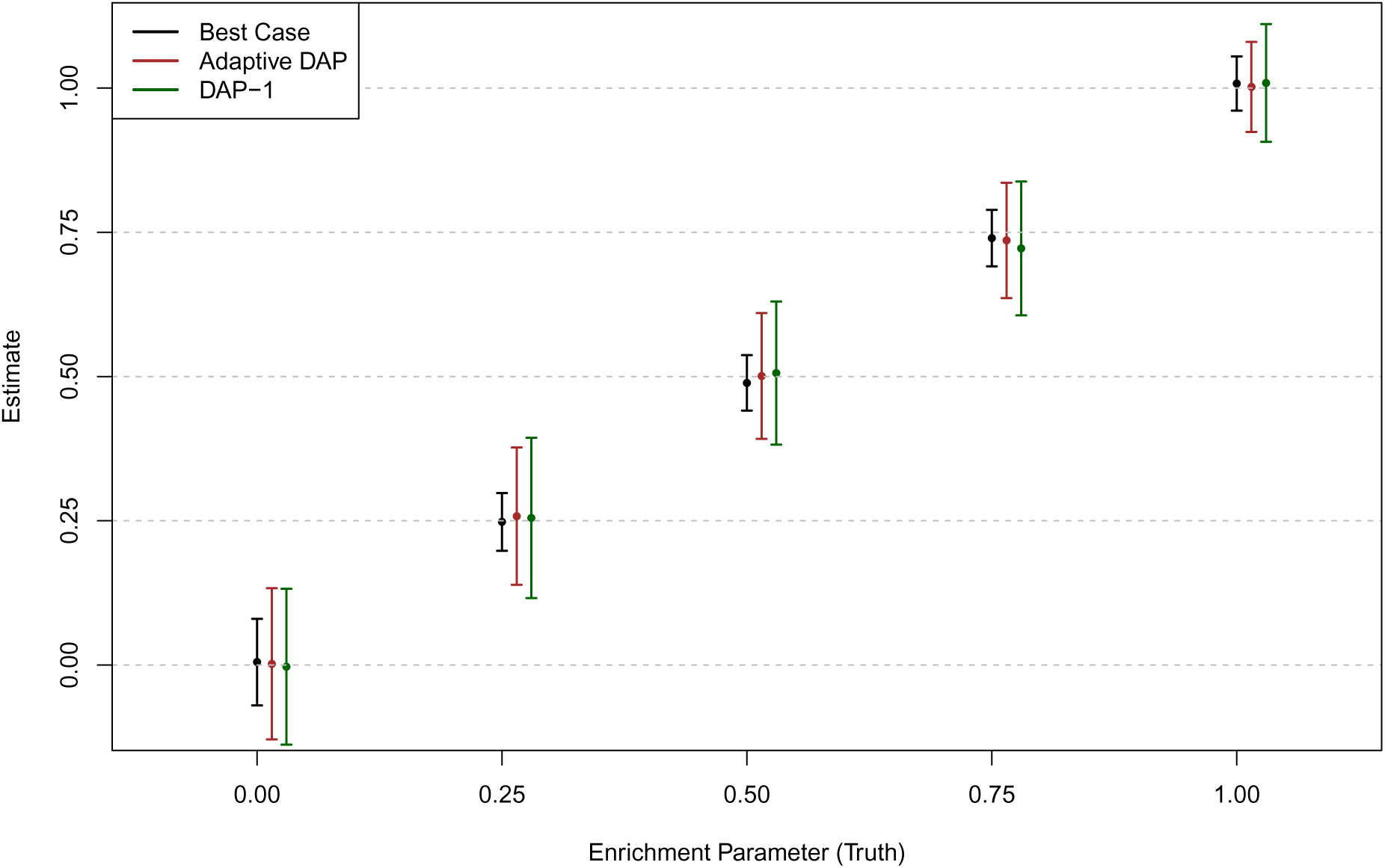
Point estimates of the enrichment parameter produced using various analysis methods in different simulation settings.

**Figure 3:**
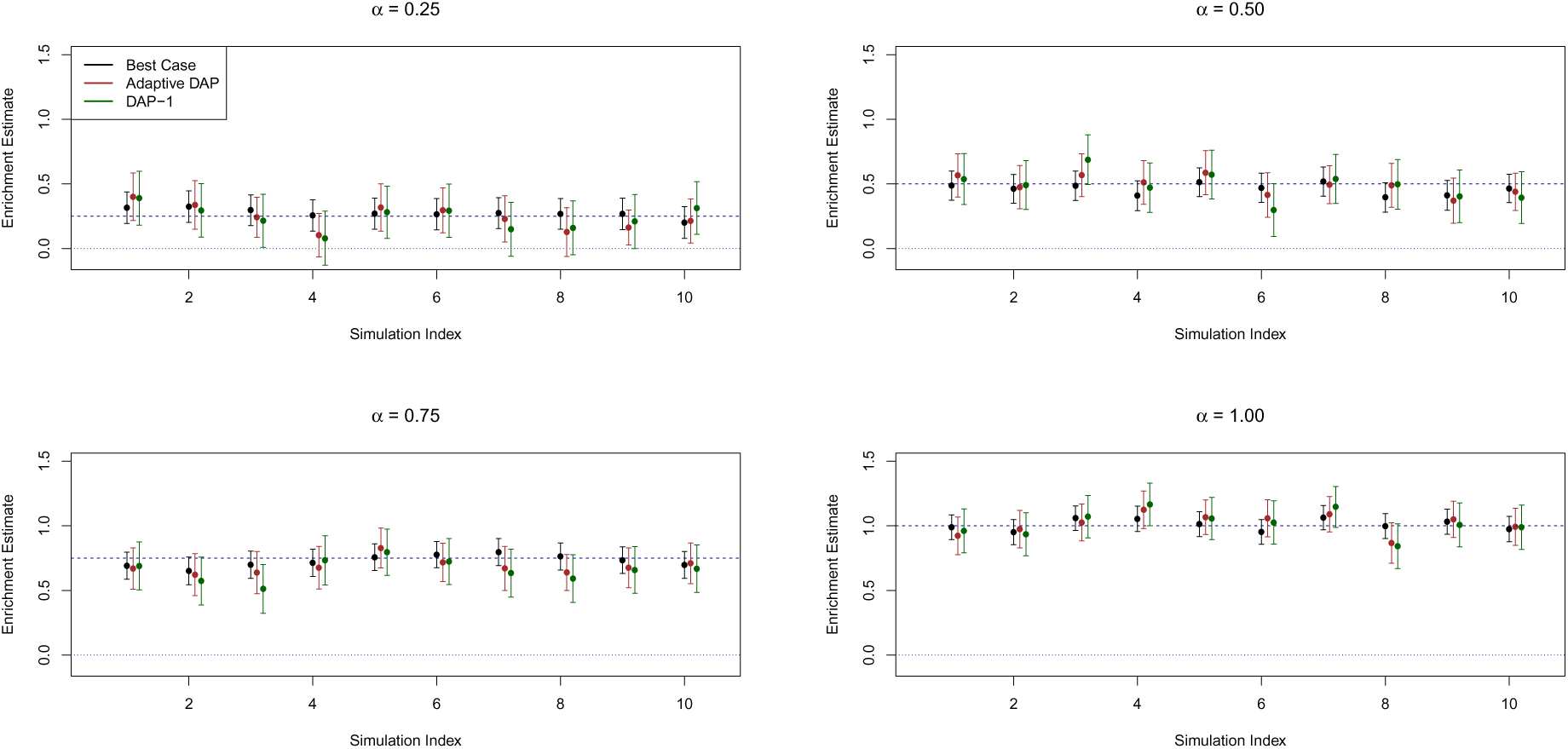
Comparison of individual estimates of the enrichment parameter and their uncertainty quantification.

The point estimate of the *α*_1_ ± standard error (obtained from 100 simulated data sets) for each method is plotted for each simulation setting. The “best case” method uses the true association status and represents the optimal performance for any enrichment analysis method. Both the adaptive DAP and DAP-1 methods yield unbiased estimates in all settings, although the adaptive DAP-embedded EM algorithm generates slightly smaller standard errors.

Each panel represents a different simulation setting. We plot the point estimates of *α*_1_ along with their 95% confidence intervals for each method using 10 randomly selected simulated data sets. In all settings, all the methods compared (“best case”, EM with adaptive DAP and EM with DAP-1) show the desired coverage probability. The figure also highlights the considerable uncertainty in enrichment analysis.

Finally, we note that both the adaptive DAP and DAP-1 algorithms underestimate the *α*_0_ parameter: on average, DAP-1 estimates *α*̂_0_ = –4.62, and the adaptive DAP yields *α*̂_0_ = –4.32 (recall that the truth is *α*_0_ = –4.00). This is fully expected, largely because of the limitation of the statistical power in detecting weak association signals. The practical consequence is that the empirical Bayes priors constructed for the final stage of multi-SNP fine mapping analysis are slightly conservative. However, we argue that the conservative priors generally lead to reduced false discoveries and may be welcomed in practice for fine-mapping analysis.

#### Accuracy of the Adaptive DAP Algorithm

In the second numerical experiment, we compare the performance of the adaptive DAP algorithm with the *exact* Bayesian computation. In particular, we are interested in evaluating the accuracy of the approximation 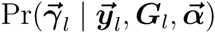 and the induced SNP-level PIP values from the adaptive DAP algorithm. The simulation setting mimics multi-SNP fine-mapping analysis at the final stage of our proposed inference procedure.

For the exact Bayesian computation with reasonable computational cost, we have to limit the number of candidate SNPs in a locus. Specifically, in each simulation, we randomly select genotypes of *p* = 15 neighboring *cis*-SNPs of a gene from the GEUVADIS data set. We then uniformly select 1 to 5 QTNs and generate the phenotype measure using a multiple linear regression model.

We apply both the adaptive DAP algorithm and the exact Bayesian posterior computation on a total of 1,250 simulated data sets using the identical prior specification. The exact computation evaluates all 2^15^ = 32, 768 association models for each simulated data set. We apply the adaptive DAP algorithm by varying the threshold value for selecting high-priority candidate SNPs, *λ*, from 0.01 to 0.05.

First, we compare the true normalizing constant *C* with the estimated value *C*\* from the adaptive DAP by computing the ratio *C*\*/*C* in each simulated data set. Utilizing all SNPs of all the simulated data sets, we also calculate the root-mean-square error (RMSE) to characterize the precision of the PIP approximations. The results indicate that for stringent *λ* values, the DAP can indeed estimate the normalizing constant with very high accuracy (Table 1 and Figure 4), which ensures the high precision of the estimated PIPs. As the *λ* threshold is relaxed, the approximation of *C* becomes less accurate in some cases; nevertheless, we observe that the overall precision level of the approximate PIPs is still reasonably high.

**Table 1:** Numerical comparison of the exact calculation and the adaptive DAP algorithm at different threshold values in the second simulation study

| $\lambda$ | Mean of $C^*/C$ | RMSE of approximate PIP |
| --- | --- | --- |
| 0.01 | 0.994 | $2.36 \times 10^{-3}$ |
| 0.02 | 0.986 | $5.32 \times 10^{-3}$ |
| 0.03 | 0.963 | $9.83 \times 10^{-3}$ |
| 0.04 | 0.921 | $1.40 \times 10^{-2}$ |
| 0.05 | 0.854 | $2.42 \times 10^{-2}$ |

**Figure 4:**
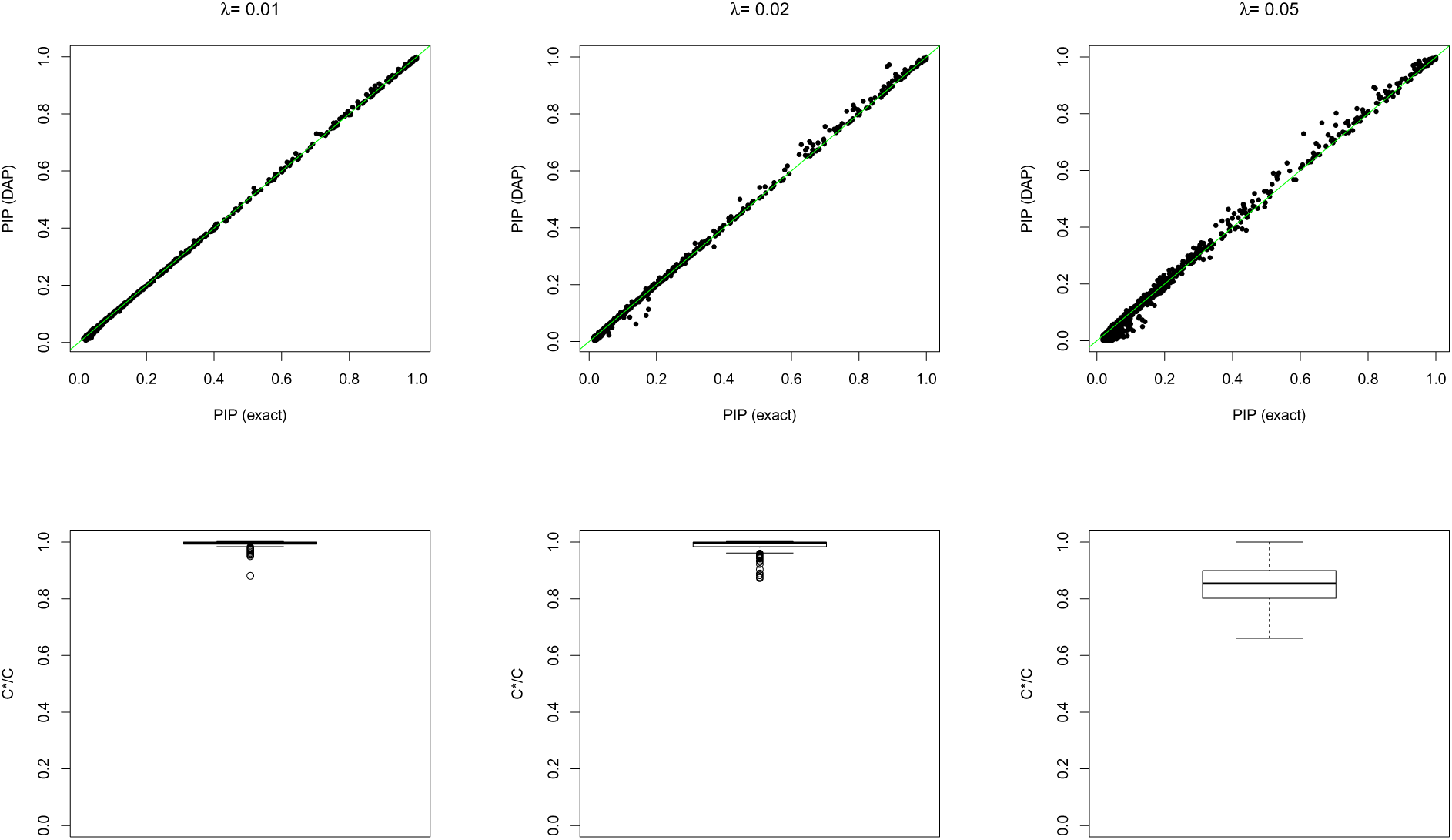
Assessment of the accuracy of the adaptive DAP algorithm at different threshold values.

In the top panel, the individual PIP approximations from the DAP are compared to the exact calculations. In the bottom panel, the distribution of *C*\*/*C* is plotted. The simulation results are obtained for threshold values *λ* = 0.01, 0.02, 0.05 for the DAP algorithm.

Next, we examine the derived stopping rule and the analytic estimation of the approximation error. Overall, we find that the stopping rule and the error approximation work extremely well for these simulations, and we summarize the results in Figure 5.

**Figure 5:**
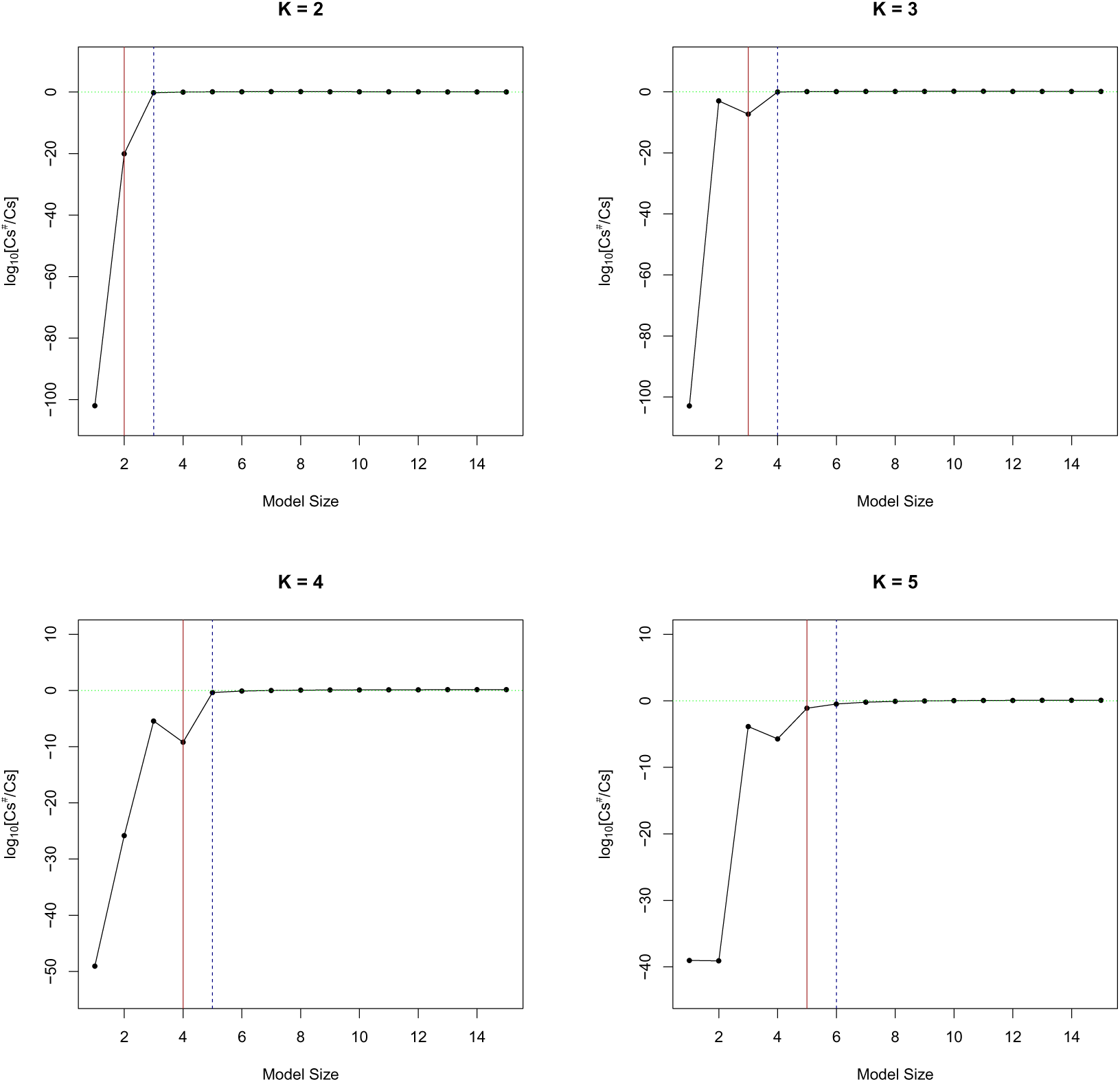
Examination of the recursive approximation of *C_s_* by equation (B.4) in the simulated data sets.

Each panel represents a simulated data set containing *K* true QTLs. The ratio of the estimated value 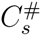 (computed using the true value of *C*_*s*–1_) over the true value *C_s_* is plotted on a log 10 scale for all model size partitions. The red vertical line indicates the size of the true association model, and the blue dotted line represents the actual stopping point at which the adaptive DAP halts explicit exploration. As the model size *s* exceeds *K*, the estimation by 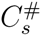 becomes very accurate in all settings.

Using the simulated data set, we also benchmark the average computational time for each simulation/analysis setting and present the results in Table 2. All runs are performed with 10 parallel threads using the OpenMP library. For the exact calculation, the average time remains constant regardless of the number of true QTNs. The DAP algorithm represents a much reduced computational time compared to the exact calculation. The general trend of the DAP running time is also clear (albeit a few small deviations): with an increasing number of true QTNs, the running time increases, and with more relaxed *λ* values, the running time decreases.

**Table 2:** Benchmark of the average computational time required for the DAP and exact computation

| Method | Running Time (seconds) |  |  |  |  |
| --- | --- | --- | --- | --- | --- |
|  | Number of True QTLs |  |  |  |  |
|  | 1 | 2 | 3 | 4 | 5 |
| DAP ($\lambda = 0.01$) | 0.097 (0.234) | 0.275 (1.180) | 0.733 (3.704) | 1.276 (7.140) | 2.527 (13.181) |
| DAP ($\lambda = 0.02$) | 0.093 (0.268) | 0.208 (0.776) | 0.663 (3.128) | 1.275 (6.816) | 2.368 (12.965) |
| DAP ($\lambda = 0.03$) | 0.087 (0.238) | 0.133 (0.408) | 0.252 (1.060) | 0.844 (4.644) | 1.422 (7.876) |
| DAP ($\lambda = 0.04$) | 0.063 (0.116) | 0.122 (0.312) | 0.230 (0.732) | 0.615 (3.064) | 0.571 (2.596) |
| DAP ($\lambda = 0.05$) | 0.050 (0.072) | 0.120 (0.280) | 0.139 (0.320) | 0.184 (0.448) | 0.180 (0.276) |
| Exact | 19.8 (121.4) |  |  |  |  |

The running time is measured in seconds by the UNIX utility program “time”. In each cell, we show the actual running time (“real” time), which is greatly reduced by parallel processing with 10 threads; in the parentheses, the “user” time is reported, which objectively reflects the actual computational cost, i.e., this measurement is not reduced by the parallelization.

#### Power Comparison of the Multi-SNP Analysis Algorithms

In the final simulation study, we compare the performance of the adaptive DAP with other existing algorithms in identifying multiple association signals. Specifically, we directly use the simulated multiple-population eQTL data sets from Wen *et al*.,^13^ where a genomic locus consisting of 100 relatively independent LD blocks (with 25 neighboring SNPs per block) is artificially assembled using real genotype data from the GEUVADIS project and 1 to 4 QTNs are randomly assigned to different LD blocks per simulation.

In Wen *et al*.,^13^ we compared three competing approaches, i) a single SNP analysis method, ii) a conditional analysis method, and iii) a multi-SNP analysis method based on an MCMC algorithm, regarding their abilities to correctly identify the QTN-harboring LD blocks. We run the adaptive DAP algorithm on the simulated data sets and compare the results with the three existing methods. Our results indicate that the adaptive DAP algorithm presents a significant improvement in performance (Figure 6) and a remarkable reduction in computational time compared with the MCMC algorithm (Table 3), and both approaches outperform the single SNP analysis and conditional analysis approaches. In addition, Figure 6 also shows that with prolonged sampling steps, the MCMC outputs seemingly “converge” to the DAP results. We also run a fast version of the adaptive DAP algorithm with tuning parameter *λ* = 0.05 (Figure 7), and the results indicate that the decrease in performance from the default setting (*λ* = 0.01) is minimum.

**Figure 6:**
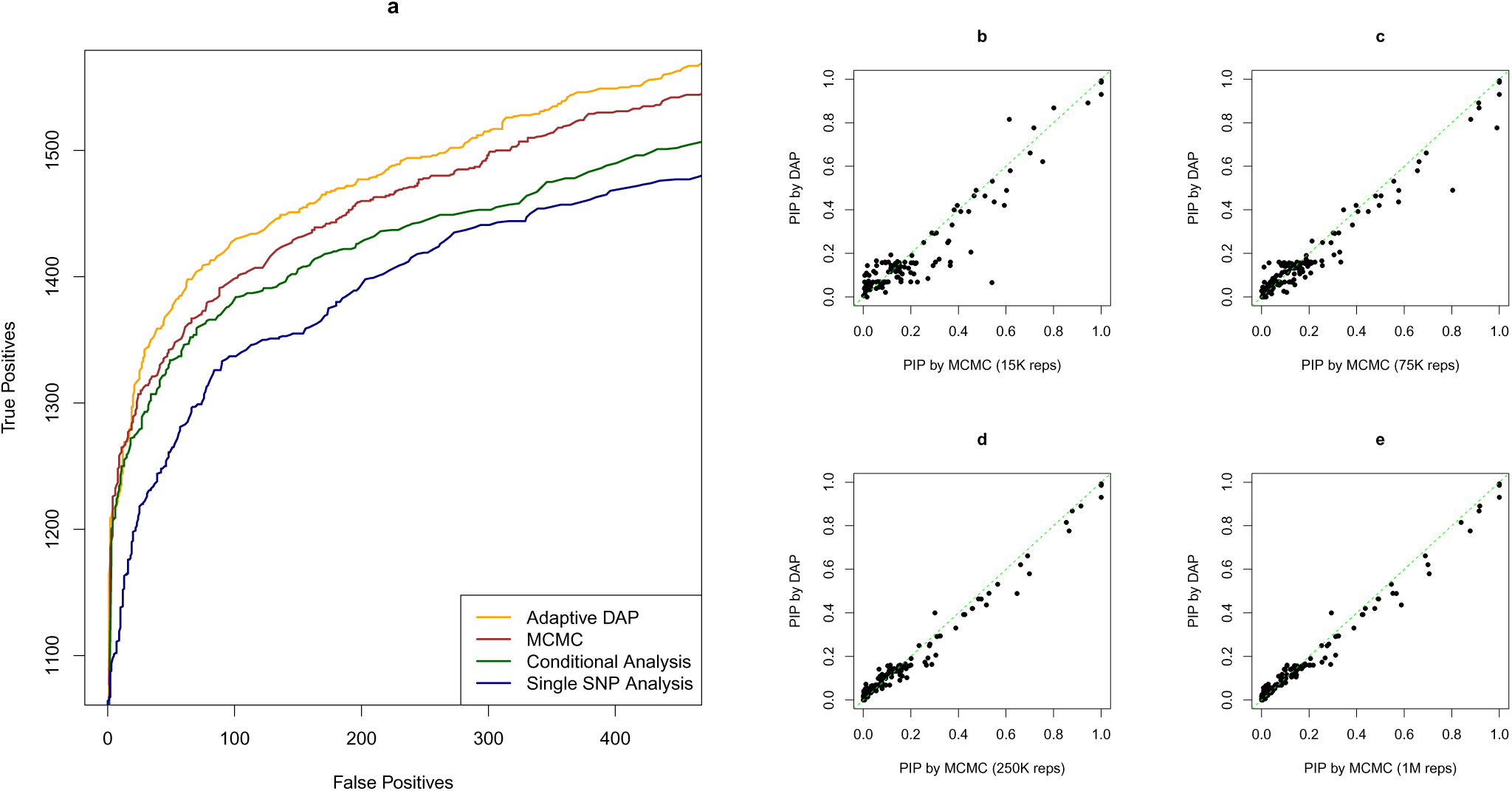
Comparison of DAP and MCMC algorithms in simulation study III. (a) Performance comparisons for multi-SNP QTL mapping. We apply different analytical approaches to a simulated data set reported in Wen *et al*.^13^ to evaluate their abilities to identify multiple independent LD blocks harboring true QTLs. The methods compared include a single-SNP analysis approach (navy blue line), a forward selection-based conditional analysis approach, the MCMC algorithm described in Wen *et al*.,^13^ and the DAP algorithm. Each plotted point represents the number of true positive findings (of LD blocks) versus the false positives obtained by a given method at a specific threshold. The MCMC algorithm and the DAP algorithm are based on the Bayesian hierarchical model and clearly outperform the other two commonly applied approaches. Most importantly, the DAP algorithm presents a significant performance improvement compared with the MCMC in both accuracy and computational efficiency. (c) - (e) Comparison of PIP values estimated by adaptive DAP and MCMC with various running lengths. We randomly selected 10 simulated data sets and ran MCMC with 4 different lengths of sampling steps, ranging from 15,000 to 1 million (the results shown in panel (a) are based on 75,000 sampling steps for each data set). With the prolonged MCMC runs, the MCMC outcomes seemingly “converge” to the DAP results.

**Table 3:** Average running time and PIP comparison using MCMC runs with varying sampling steps in simulation study III. The actual running time reported from the UNIX “time” command is shown for each experiment. The DAP algorithm runs with 10 parallel threads, and the average user time (i.e., approximate running time without parallelization) is 1 minute and 8.66 seconds.

| | MCMC (reps) | | | | DAP<br>$\lambda = 0.01$ |
| --- | --- | --- | --- | --- | --- |
|  | 15K | 75K | 250K | 1M |  |
| Running Time (real) | 4m 2.79s | 10m 28.37s | 28m 50.00s | 107m 46.75s | 28.44s |
| RMSE of PIP (w.r.t DAP) | 0.080 | 0.052 | 0.034 | 0.030 | — |

**Figure 7:**
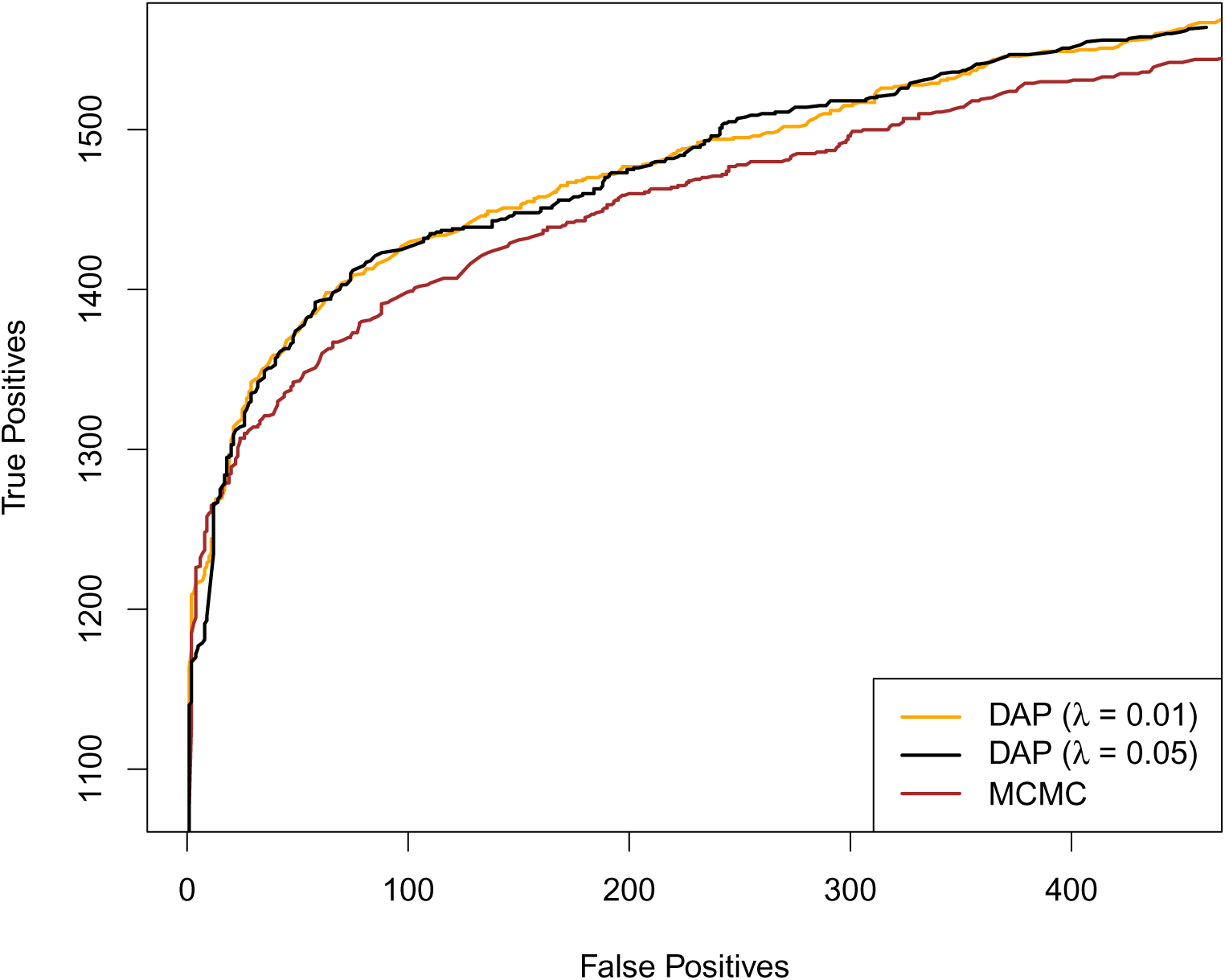
Additional comparisons for multi-SNP QTL mapping.

We show the additional simulation results by running the adaptive DAP with *λ* = 0.05, which is most similar to the DAP outcome with the default setting (*λ* = 0.01) and, for the most part, still outperforms the MCMC algorithm.

### Re-analysis of the GEUVADIS Data

We re-analyze the cross-population eQTL data set generated from the GEUVADIS project (Web Resources) using the proposed 3-stage inference procedure. In this re-analysis, we focus on examining two types of genomic annotations that are known to impact the enrichment of eQTNs: the SNP distance to the transcription start site (TSS) of the target gene and annotations assessing the ability of a point mutation to disrupt transcription factor (TF) binding. Following Wen *et al*.,^13^ we group all SNPs within 100 kb of a gene into 1 kb non-overlapping bins according to their distances from the TSS and use the label of the corresponding bin for each SNP to represent its distance to TSS (DTSS) as a categorical variable. In addition, a SNP is classified as a *binding SNP* if it is computationally predicted to strongly disrupt TF binding by the CENTIPEDE model using the ENCODE DNaseI data^26,27^ (Web Resources). If a SNP is located in a DNaseI footprint region but there is no strong evidence for disrupting TF binding, it is classified as a *footprint SNP*; otherwise, the SNP is labeled as a *baseline SNP.* Due to the computational restraint, our previous enrichment analysis reported in Wen *et al*.^13^ was based on a single iteration of the MCMC-within-EM (or EM-MCMC) algorithm (i.e., the E-step is carried out by the MCMC algorithm), as our main goal was enrichment *testing*. Although the evidence is sufficiently strong for testing purposes, the enrichment parameters were known to be severely underestimated.

We ran the complete DAP-1-embedded EM algorithm to perform the enrichment analysis. The full EM algorithm runs for 25 iterations to meet our convergence criteria, which require an increment ≤ 0.01 in the log-likelihood between two consecutive iterations (Figure S5). The complete EM run takes 21 minutes on a Linux box with a single 8-core Intel Xeon 2.13 GHz CPU. In comparison, the MCMC algorithm takes approximately 84 hours of computational time to fully process all 11,838 genes in a *single* E-step on the same computing system.

After a single iteration, the DAP-1-embedded EM algorithm yields point estimates for the TF binding annotations that are very similar to our previous results reported in Wen *et al*.^13^ (Table 4). As expected, the final estimates from the complete EM run have very high enrichment values: the binding SNPs have an estimated log odds ratio *α*̂_1_ = 0.94, or fold change of 2.56, with the 95% CI [0.84,1.05], whereas the footprint SNPs have a much lower enrichment estimate (log odds ratio *α*̂_1_ = 0.53 or fold change of 1.70, with the 95% CI [0.40, 0.67]). Note that the two confidence intervals are non-overlapping. In comparison, our previously reported estimates of the corresponding enrichment parameters are 0.40 (95% CI [0.32, 0.49]) and 0.14 (95% CI [0.04, 0.24]) for binding and footprint SNPs, respectively.

**Table 4:** Comparison of enrichment estimates by EM-DAP1 and EM-MCMC after a single iteration in analysis of GEUVADIS data

| Method | Footprint SNPs |  | Binding Variants |  |
| --- | --- | --- | --- | --- |
| | $\alpha$ | 95% C.I. | $\alpha$ | 95% C.I. |
| EM-MCMC | 0.14 | (0.04, 0.24) | 0.39 | (0.32, 0.49) |
| EM-DAP1 | 0.12 | (−0.01, 0.25) | 0.41 | (0.30, 0.51) |

The binding SNPs refer to the genetic variants that are computationally predicted to disrupt TF binding, and the footprint SNPs are those simply located in the DNaseI footprint region but not predicted to affect TF binding. The enrichment estimates from both methods are very similar. The MCMC algorithm accounts for multiple independent association signals and yields slightly tighter confidence intervals, as expected. However, the EM-DAP1 is much more computationally efficient: it runs almost one thousand times faster than the EM-MCMC algorithm.

Next, we repeat the multi-SNP fine-mapping analysis using the adaptive DAP algorithm and the new set of the empirical Bayes priors obtained from the enrichment analysis. For most genes, the results (i.e., the number of independent signals for each gene) are qualitatively unchanged compared to the previous MCMC results. Nevertheless, we find that fine-mapping with the adaptive DAP is much more efficient, and the annotated SNPs, especially the binding SNPs, are further prioritized in the new fine-mapping results (Figure 9).

**Figure 8:**
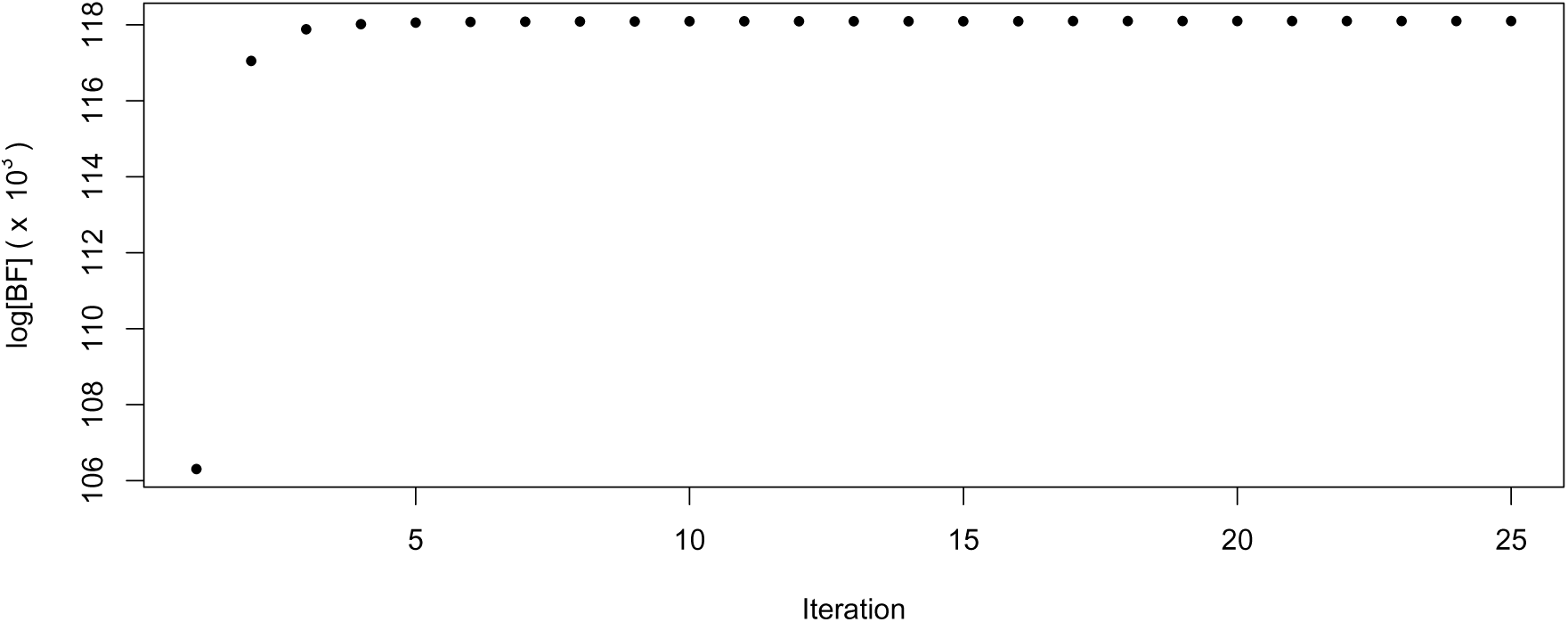
Traceplots of the marginal likelihood (in Bayes factor on the log scale) during the DAP-1-embedded EM run for analyzing the GEUVADIS data.

**Figure 9:**
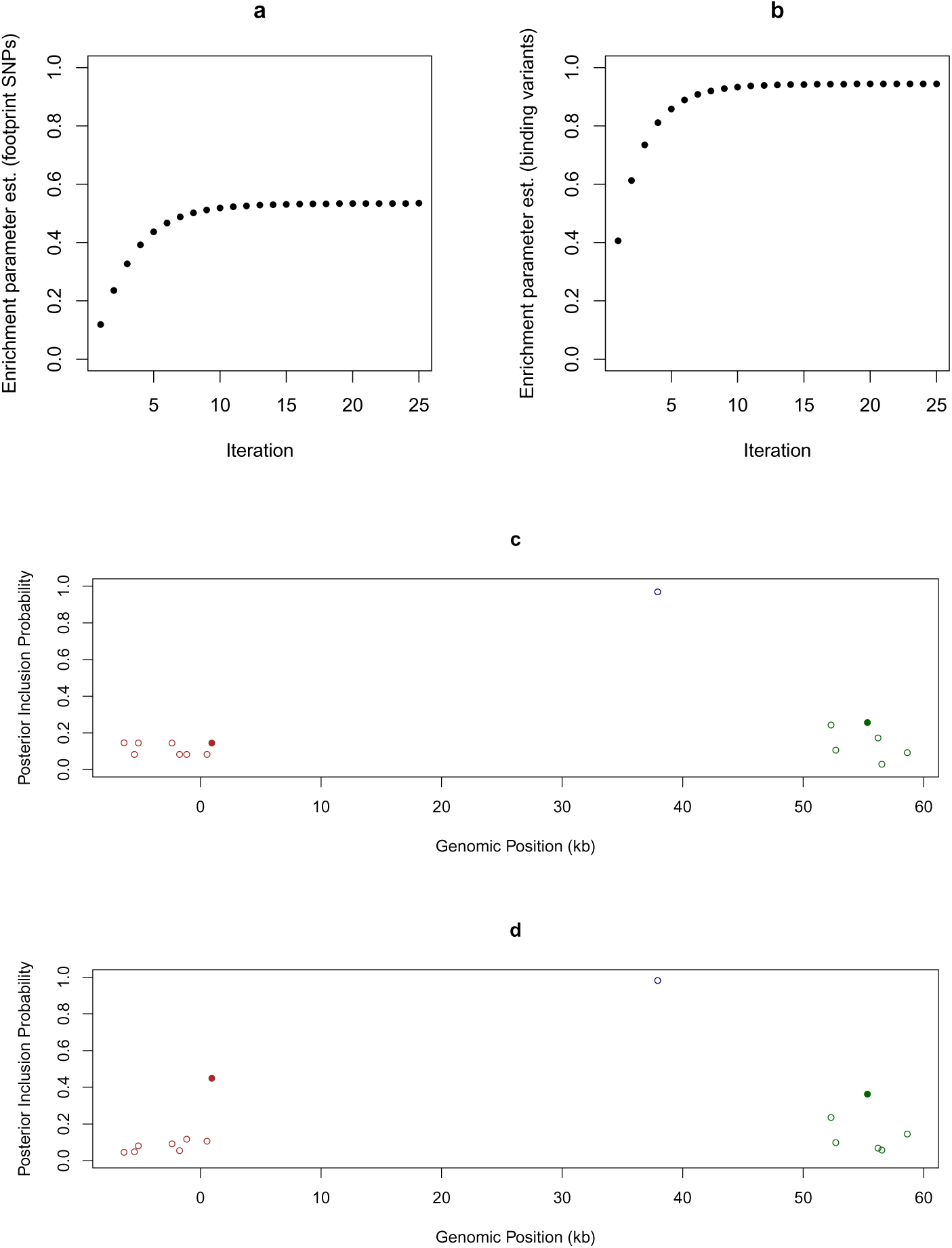
Figure 9: Output from the re-analysis of GEUVADIS data. (a) - (b) Traceplots of estimates of the enrichment parameters for binding variants and footprint SNPs during the DAP-1-embedded EM iterations for analyzing the GEUVADIS data. Both estimates are stabilized after approximately 8 iterations. (c) - (d) Comparison of multi-SNP *cis*-eQTL mapping with and without incorporating functional annotations. We plot the multi-SNP QTL mapping results of *LY86* [MIM 605241] using the GEUVADIS data. Panel (c) shows the results assuming that all SNPs are equally likely to be associated *a priori*, i.e., no functional annotation is used. Panel (d) shows the results using the functional annotations with enrichment parameters estimated by the DAP-1-embedded EM algorithm. In both cases, we use the adaptive DAP algorithm to perform the multi-SNP QTL mapping and plot the SNPs with PIP > 0.02 with respect to their positions relative to the transcription start site. SNPs in high LD are plotted with the same color, and the filled circles indicate that a SNP is annotated as disrupting TF binding. It is clear that three independent *cis*-eQTLs exist because in both panels, the sums of the PIPs from the SNPs with the same color all → 1. When incorporating functional annotation to perform integrative QTL mapping, the binding variants show much greater PIP values and are prioritized over the non-annotated SNPs in high LD.

### Analysis of the GTEx Data

We analyze the *cis*-eQTL data from the GTEx project (Web Resources). One of the most unique advantages of the GTEx data is that they enable the study of the commonality and specificity of the eQTLs in multiple tissues. Taking advantage of the high computational efficiency of the EM-DAP1 algorithm, we perform the enrichment analysis of the TF binding annotations, derived from the ENCODE data and the CENTIPEDE model, in eQTLs across 44 human tissues while controlling for the SNP distance to TSS. More specifically, for each gene, we consider a 2 Mb *cis* region centered at the transcription start site. For each tissue, we perform the enrichment analysis using two sets of TF binding annotations, one derived from the ENCODE LCL cell-line and the other from the ENCODE liver-related HepG2 cell-line^27^ (Web Resources). This exercise aims to assess the impact of the cell type-specific annotations on the proposed integrative analysis.

Our results indicate that the binding variants are significantly enriched in eQTLs in all tissues regardless of the origin of the annotations. Furthermore, the point estimates of enrichment levels for binding variants are consistently higher than those for footprint SNPs, except in one occasion (small intestine tissue with LCL-derived annotations) where the two estimates are indistinguishable. Importantly, we find that the enrichment estimates in specific tissues are quantitatively correlated with the origins of the annotations. Figure 10 shows the results of the enrichment level estimates (*α*̂_1_) of the binding variants in each tissue using the LCL- and HepG2-derived TF binding annotations. Most interestingly, the LCL-derived annotations yield the highest enrichment estimates in LCLs and whole blood from the GTEx data sets, whereas the liver-related HepG2-derived annotations obtain the highest enrichment estimate in the GTEx liver tissue. Overall, our results suggest that TF binding annotations derived from different tissues must have substantial overlaps; nevertheless, the annotations from the relevant tissues may provide better functional interpretations for expression-altering causal SNPs in a specific tissue.

**Figure 10:**
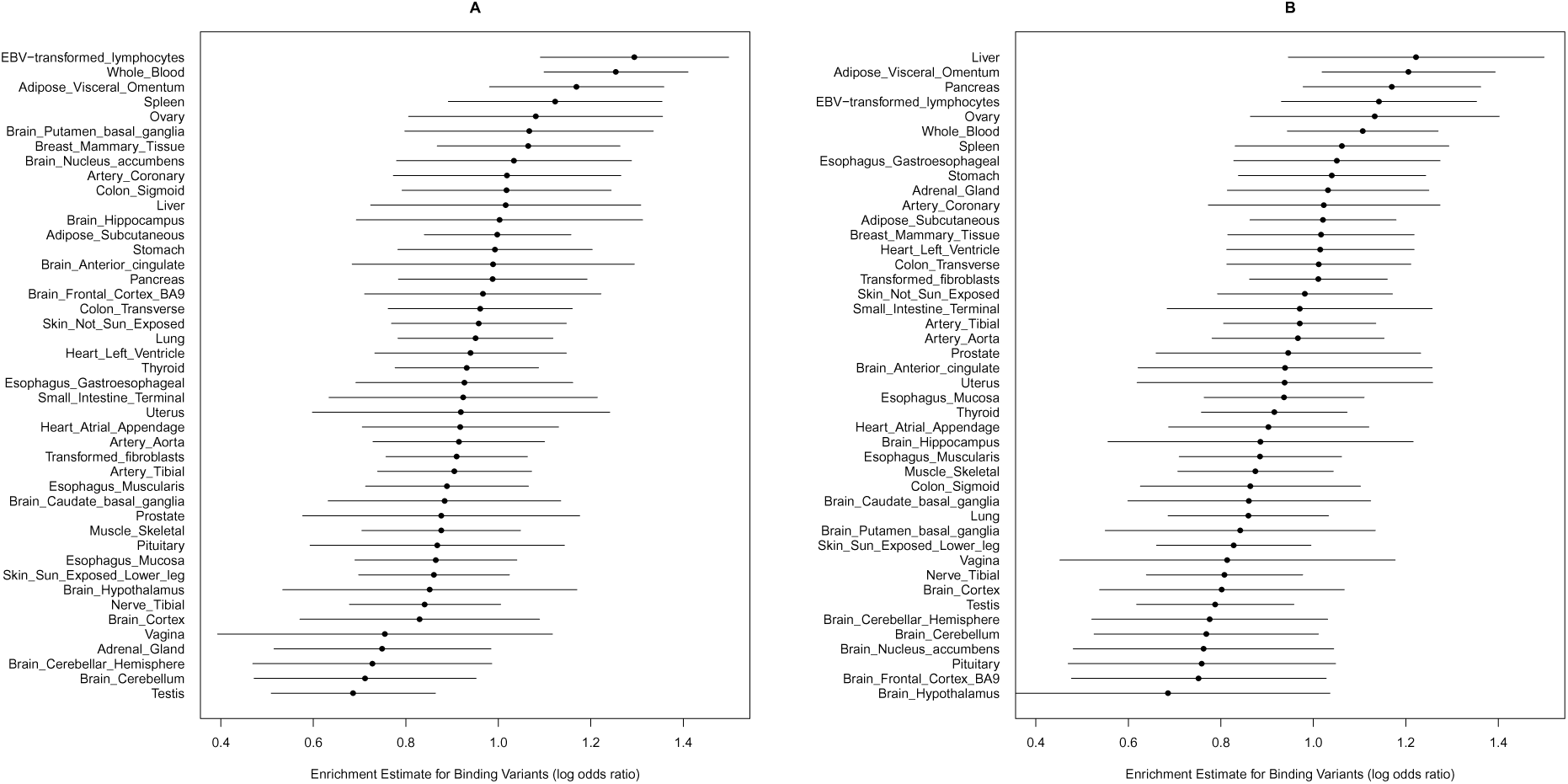
Enrichment estimates for binding variants in GTEx tissues.

The estimates in panel A are based on the annotations derived from the DNaseI data of the ENCODE LCLs, whereas the estimates in panel B are based on annotations derived from the ENCODE liver-related HepG2 DNaseI data. In each panel, we plot the point estimate of the enrichment parameter and its 95% confidence interval in each tissue. The tissues are ranked in descending order according to the magnitude of the point estimates. All estimates are obtained controlling for the SNP distance from TSS. All estimates are significantly far from 0 (at the 5% level). Interestingly, when the tissue and origin of the annotations match, the point estimates for enrichment are the highest.

The top, middle and bottom panels display the histogram of the posterior expected number of *cis*-eQTLs from all the eGenes in the liver, lung and blood tissues, respectively. For most genes, we can only identify a single association signal. However, for a non-trivial number of eGenes, multiple independent association signals can be confidently identified by the adaptive DAP algorithm. The sample size is seemingly an important factor related to the ability to identify multiple independent signals in a *cis* regfion.

We then proceed to identify genes that harbor QTNs (i.e., eGenes) using a Bayesian FDR control procedure that we recently developed.14 Subsequently, we perform multi-SNP fine-mapping analysis for the identified eGenes incorporating the enrichment estimates using the adaptive DAP algorithm. We present the analysis results for the liver (sample size 97), lung (sample size 278) and whole blood (sample size 338). There are 2,788, 8,605 and 7,937 eGenes that are identified from the lung, liver and whole blood, respectively. We suspect that the number of differences in eGenes discovery is largely attributed to the sample sizes but is also correlated with the levels of experimental noise in measuring the gene expression in each tissue. For each fine-mapped eGene *l* in each tissue, we compute the posterior expected number of independent signals using 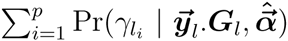 and plot the histogram for each tissue in Figure 11. In all three tissues, we identify single eQTL signals for the vast majority of eGenes. Nonetheless, for a non-trivial number of genes, we are able to confidently identify multiple independent signals. Comparing the fine-mapping results among the three tissues, we find that the ability to identify additional independent signals is also seemingly correlated with the sample sizes.

**Figure 11:**
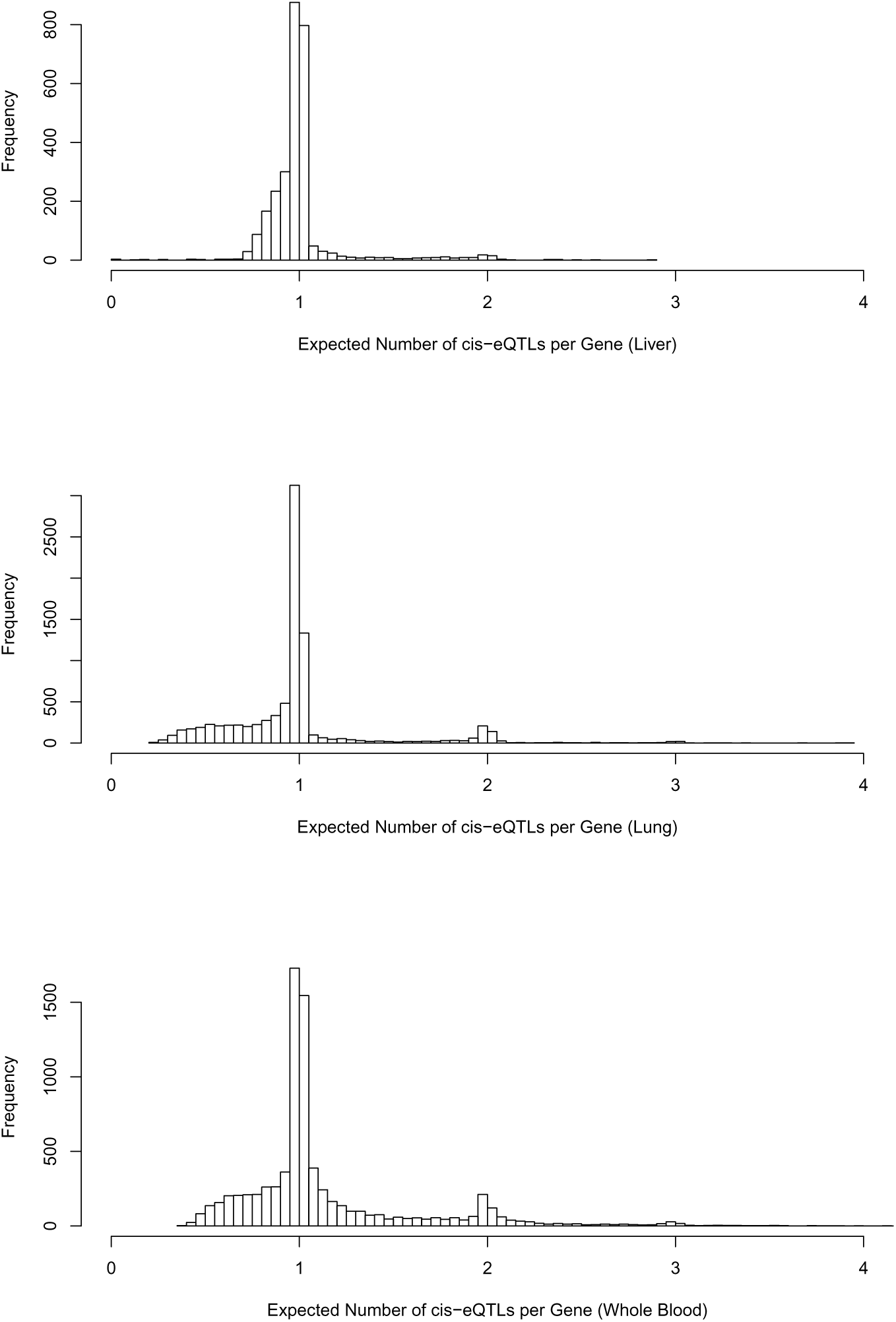
Posterior expected number of *cis*-eQTL signals per eGene in GTEx liver, lung and whole blood tissues.

We further examine some known individual genes to validate our integrative analysis results. In particular, we examine *SORT1* [MIM 602458], whose function is related to plasma low-density lipoprotein cholesterol (LDL-C [MIM 613589]) metabolism through modulation of hepatic VLDL secretion. Through GWAS meta-analysis and extensive functional analysis,^28^ a single SNP, rs12740374, is identified to cause variations in LDL-C. More specifically, the major allele disrupts the binding site of C/EBP transcription factors in human hepatocytes. Our integrative fine-mapping analysis using the GTEx liver data yields a Bayesian 95% credible set, narrowed down to only two potential causal eQTNs for *SORT1*: rs12740374 (PIP = 0.473) ranks second very closely only to SNP rs7528419 (PIP = 0.526). Moreover, the direction of the genetic effect for rs7528419 fits the description provided in Musunuru *et al*.^28^ The two SNPs in the credible set are in high LD (*r*^2^ > 0.95), except that the genotypes of rs12740374 in the GTEx samples are *not* directly genotyped but imputed. Upon further investigation, we find that the binding site reported by Musunuru *et al*.^28^ is not captured by the ENCODE DNaseI experiments in HepG2, and hence, rs12740374 is not correctly annotated. We then include the annotation of rs12740374 as a binding SNP based on the functional study of Musunuru *et al*.^28^ and re-run the fine-mapping analysis using the adaptive DAP. We find that rs12740374 yields the highest PIP value (PIP = 0.752) among all the candidate SNPs (the PIP for rs7528419 drops to 0.247). The lesson learned here is that the completion of the genomic annotations may have a profound impact on the integrative analysis, and efforts should be made to generate a more comprehensive set of genomic annotations by both accumulating new experimental data and integrating them with all the existing data.

## Discussion

The proposed EM-DAP1 algorithm provides an efficient and flexible framework to perform enrichment analysis with respect to genomic annotations using genetic association data – there is no restriction on the types of annotations (categorical or continuous) or the number of annotations that can be simultaneously investigated. Some of the commonly applied *ad-hoc* enrichment analysis methods in the same context attempt to first classify the binary latent association status **Γ** for all candidate SNPs based on their single SNP testing results. However, it is worth noting that the classification based on hypothesis testing typically has very stringent controls over type I errors but is much more tolerant (in practice, it may be too tolerant) and has little control over type II errors, which are a major source of the overall mis-classification errors for **Γ**.^13^ As a consequence, most *ad-hoc* procedures of this type provide poor quantification of enrichment levels. Recently, probabilistic model-based enrichment analysis approaches have been proposed based on the “one QTN per locus” assumption and applied to both molecular QTL mapping and GWAS.^11^ A common feature of these approaches is that they treat each locus as the exchangeable/comparable unit in the analysis: in the simplest case, each locus has the common prior probability, *π*_1_, of harboring causal QTNs. Although the DAP-1 algorithm implicitly also makes the same assumption and enjoys the benefit of fast and efficient computation using only summary statistics, it presents some significant differences/improvements compared to the aforementioned approaches. The DAP-1 algorithm, built on the proposed hierarchical model, considers each SNP as the unit of analysis. This modeling strategy leads to a straightforward EM algorithm for parameter estimation, where the target function in the M-step is convex with well-known optimization solutions. In comparison, with the parameterization including *π*_1_, the target function in the M-step is no longer guaranteed to be convex, which can cause convergence issues in EM estimation and prevent the simultaneous investigations of many annotations (see the details in the Appendix C). Furthermore, *π*_1_ parameterization essentially assumes that genetic loci consisting of many SNPs are equally likely to harbor causal QTNs as loci consisting of only a few SNPs. From the empirical evidence produced by eQTL analysis, we find that this assumption is likely false : the genes with more *cis* candidate SNPs are more likely to harbor eQTNs.^13^ In summary, the proposed hierarchical model and the EM-DAP1 algorithm represent better alternatives.

The proposed Bayesian hierarchical model does not explicitly consider potential polygenic background. To evaluate the performance of the proposed enrichment analysis method under an explicit polygenic model, we modify the simulation settings for enrichment analysis by imposing a small yet non-zero genetic effect on every candidate SNP. Under such setting, *γ_i_* should be interpreted as an indicator whether the genetic effect of SNP *i* is significantly larger than the polygenic background. The simulation results (Figure 12) indicate that the estimates of the enrichment parameters are biased toward 0 in the presence of polygenic background: although the bias is negligible when the polygenic effects are small. We plan to extend our current work to fully account for polygenic background in our future work by considering a more appropriate model like the Bayesian sparse linear mixed model (BSLMM).^29^

**Figure 12:**
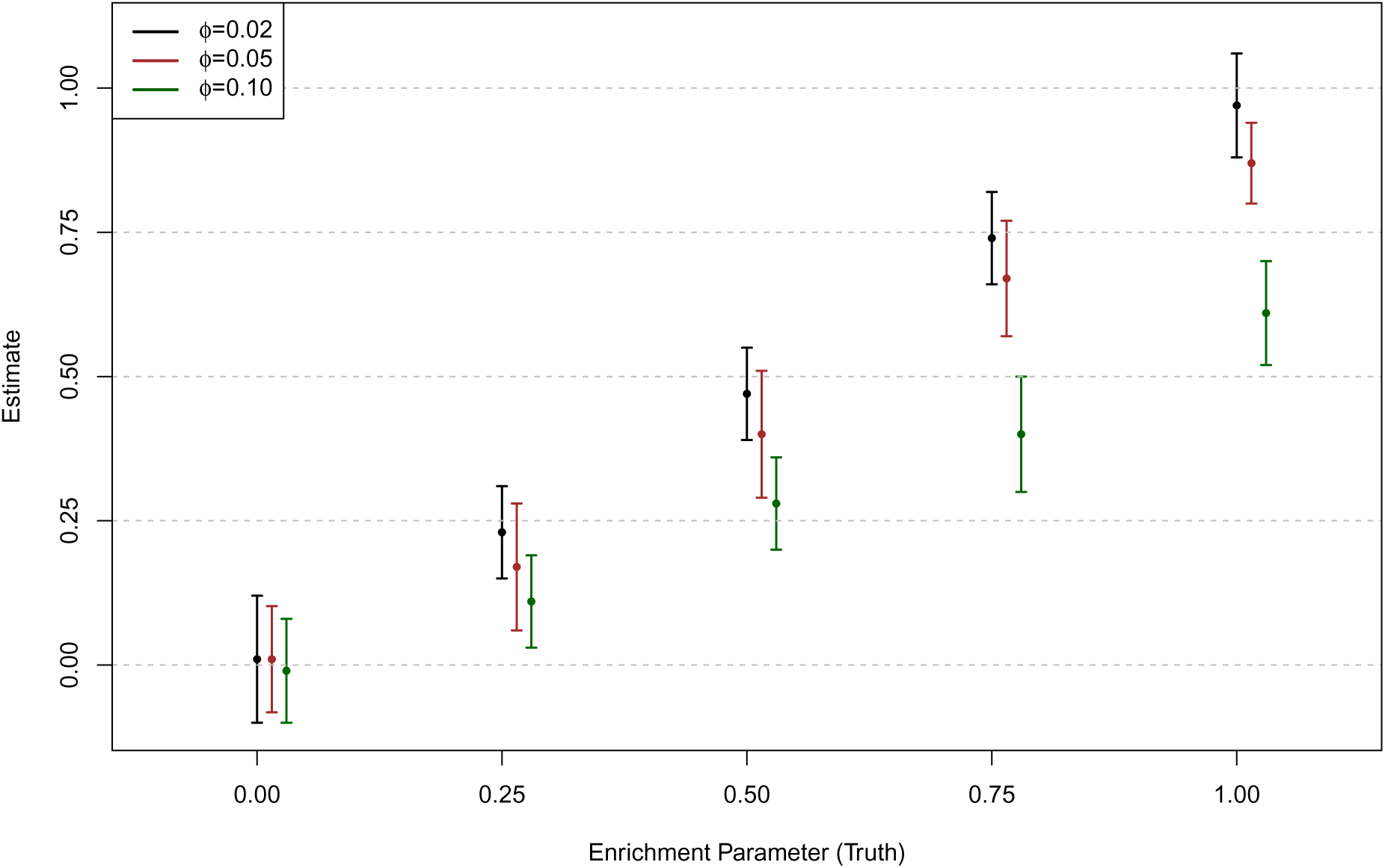
Estimates of the enrichment parameters for data simulated from polygenic models.

In this experiment, the simulation scheme is mostly similar to the first simulation study described in the main text, except that in addition to the SNPs sampled to have large effects, we assign a non-zero genetic effect from an independent N(0, *ϕ*^2^) distribution for all the remaining candidate SNPs. (In this case, *γ_i_* should be interpreted as an indicator of large genetic effect.) We select *ϕ* = 0.02, 0.05 and 0.1 to represent different magnitude of polygenic background. The point estimate of the *α*_1_ ± standard error (obtained from 50 simulated data sets using DAP-1-embedded EM algorithm) for each *ϕ* value is plotted. In all cases, the non-zero *α*_1_ estimates are biased toward 0, however when *ϕ* is small (*ϕ* = 0.02), the bias seems negligible.

Our analysis of multi-tissue eQTL data yields many interesting findings that are worthy of indepth follow-up investigation. In particular, our results suggest that the cell type specificity and the completeness/accuracy of the genomic annotations may have profound impacts on the integrative association analysis in terms of different aspects as follows: the cell-type specificity of the annotations affects the global enrichment estimates and the multi-SNP analysis results of *every* subsequently fine-mapped locus, whereas mis-annotations of certain variants likely impact functional interpretations of specific loci but are not likely to alter the global enrichment estimates as long as the annotations are accurate *on average*. These findings should motivate efforts to generate a more comprehensive and accurate catalog of genomic annotations to improve the overall quality of genetic association analysis. Furthermore, it should be noted that all the annotations could have additional levels of complexity (e.g., *cis* regulatory grammar) that can be consistently analyzed within the same framework by extending our logistic prior model in a straightforward manner to allow interactions. To aid these efforts, our proposed genome-wide scale enrichment analysis has provided a principled way of assessing the tissue/cell type specificity of the genomic annotations.

## Acknowledgments

We thank the GTEx consortium and the GEUVADIS RNA sequencing project for releasing valuable data in a timely fashion. This work is supported by NIH grants MH101825 (XW), HG007022(XW) and GM109215 (XW, YL, FL and RP).

## Web Resources

The URLs for data presented herein are as follows,

DAP software and tutorial, 

~~~
http://github.com/xqwen/dap/
~~~

GUEVADIS data, 

~~~
http://www.geuvadis.org/web/geuvadis/rnaseq-project
~~~

Re-analyzed multi-SNP fine-mapping results of the GUEVADIS data, 

~~~
http://www-personal.umich.edu/~xwen/geuvadis/new_fm_rst/
~~~

GTEx data, 

~~~
http://www.gtexportal.org/home/datasets
~~~

Transcription factor binding site annotations by the extended CENTIPEDE model, 

~~~
http://genome.grid.wayne.edu/centisnps/
~~~

## Appendices

## Appendix A Selection of Priority SNPs in Adaptive DAP

We give a detailed account of the Bayesian conditional analysis procedure for selecting high-priority SNPs in the adaptive DAP algorithm. For a given locus *l*, the procedure starts with model size partition *s* = 1. Let 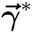 denote the model with the highest posterior probability in the size partition *s* – 1 in locus *l*, i.e.,

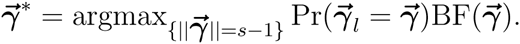

For each SNP *i* that is *not* included in the current best model, we compute a Bayes factor for the expanded model, 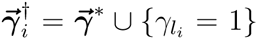. Assuming that there is exactly one additional QTL and that each candidate SNP *i* is equally likely to be *the* additional causal association *a priori*, the corresponding conditional posterior probability for SNP *i* can be computed by

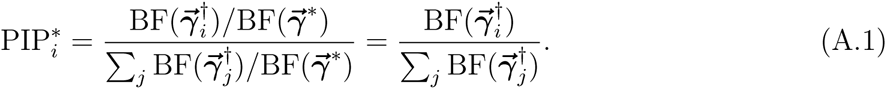

The resulting quantity is a well-defined posterior probability and is solely determined by the relative likelihood values of the expanded models. In particular, it should be noted that (A.1) fully accounts for LD between SNPs: e.g., if two SNPs are in perfect LD, they would possess identical values that correctly reflect the uncertainty (i.e., they are indistinguishable). The procedure requires *p* – *s* evaluations of Bayes factors that are computationally trivial for small *s* values. Given the pre-defined threshold *λ*, we add the SNP i into the existing set of high-priority SNPs if it is not already in the set and 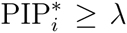. For *s* ≥ 2, we then enumerate all s-combinations from the resulting set of priority SNPs to compute 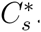. During this enumeration, we also record the new 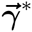 for the increased model size.

Intuitively, the threshold parameter *λ* is related to the precision of the approximate PIPs. The selection procedure roughly estimates the probability, 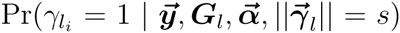, for SNP *i*. Note the relationship

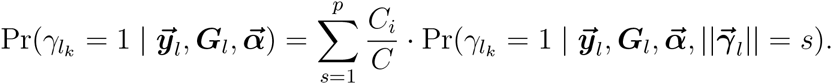

The following can be concluded:

1. If 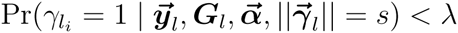 for a given SNP at all s values, then it must be the case that the overall PIP < *λ*.
2. The loss of precision of the PIP of SNP *i* due to the selection screening for a particular size partition must be < *λ*.

## Appendix B Stopping Rule and Estimation of the Approximation Error in Adaptive DAP

When a non-associated SNP is added to an existing association model, the marginal likelihood of the model is typically non-increasing. In fact, the marginal likelihood measured by the corresponding Bayes factor usually decreases slightly due to the effect of Occam’s razor built into the Bayes factor computation.^30^ We utilize this property to reduce the computation of DAP by eliminating unnecessary explicit explorations of the model partitions once the sizes of the models are considered saturated. To achieve this goal, the DAP starts the exploration with model size partition *s* = 1 for increasing s values until a stopping rule is met. The contribution of the unexplored size partitions (i.e., the approximation error) is then estimated by an analytic combinatorial approximation.

To explain the stopping rule and the combinatorial approximation, we assume that there are *K* detectable true QTNs. In each model size partition where *s* > *K*, we can classify all models into (*K* + * 1) mutually exclusive categories according to the number of true QTNs (0 to *K*) included in each association model. In the category including exactly *m* true QTLs, each member association model also includes (*s* – *m*) non-associated SNPs, and the total number of the association models in the category is given by 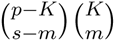. We estimate the contribution to 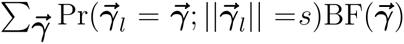 from this particular category by the equation

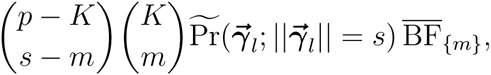

where 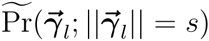 represents the average prior value within the category and 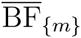 is the average Bayes factor across models including m out of K detectable QTNs. The use of 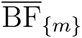 is mainly based on the assumption that including non-associated SNPs in an association model does not, on average, increase the marginal likelihood/Bayes factor. Hence, we obtain

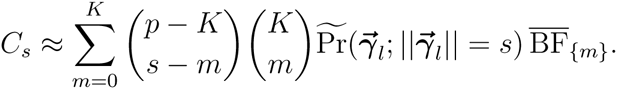

To relate *C*_*s* * 1_ to *C_s_*, we note that

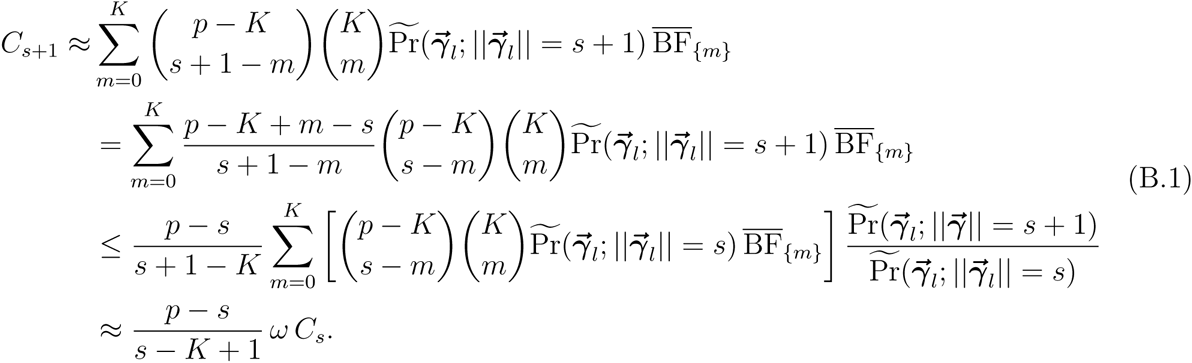

In the last step, we approximate the quantities 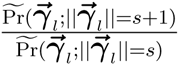 in all *K* + * 1 categories by the average prior odds 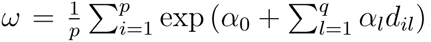. Similarly, we can derive an approximate lower bound for *C*_*s* * 1_

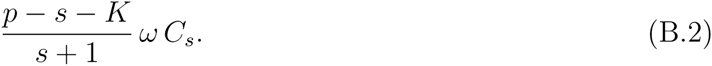

Thus, we have shown

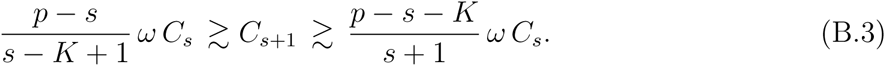

Because *K* is unknown, we estimate *C*_*s* + * 1_ from *C_s_* by the following approximation

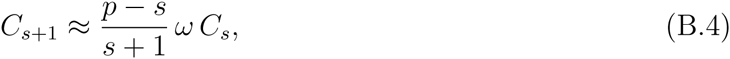

which does not depend on *K* and lies in the interval 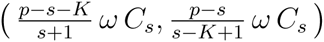. Our numerical experiment shows that this approximation is surprisingly accurate (Figure S3).

Our stopping rule is built upon the upper bound specified by the inequality (B.3). Specially, the adaptive DAP stops explicit exploration at partition size *s* = *t* if

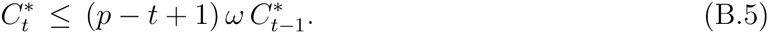

The inequality essentially tests *K* ≥ *t* – 1. In addition to utilizing the combinatorial approximation, the DAP further monitors the increment of the partial sum 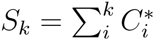. To ensure a high accuracy of the approximation, we also add an optional criterion to the stopping rule on top of (B.5), i.e.,

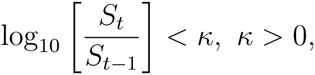

or, equivalently,

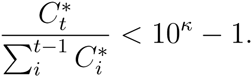

By default, we set *κ* = 0.01, which further ensures that the subsequent model size partitions make no substantial contributions to the normalizing constant. This additional criterion provides practical flexibility for running the DAP: as *κ* → 0, it enforces the DAP to explore all the model size partitions, whereas when κ is large, only the stopping rule (B.5) is effective.

Once the stopping rule is invoked, we estimate *ϵ* by

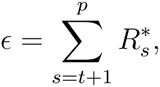

where we define 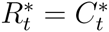 and

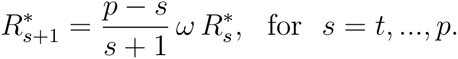

## Appendix C Derivation of the DAP-1 Algorithm

In this section, we provide a detailed derivation for the DAP-1 algorithm. It should be noted that the derivation can be generalized to the DAP-K algorithm with *K* > 1.

The key assumption of the DAP-1 is that posterior probabilities of single-QTL association models dominate the posterior probability space of 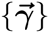, i.e.,

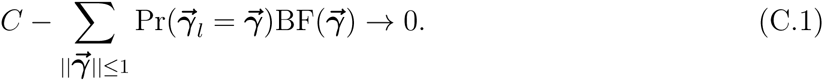

Consequently, it follows that

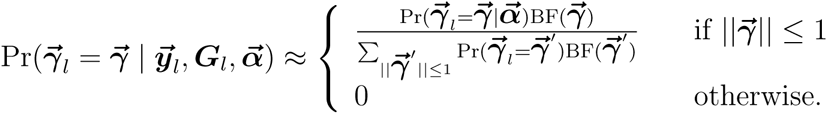

The model space of 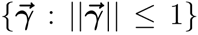 contains only the null model, 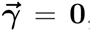, and all single-SNP association models. For the null model, it is clear that 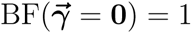, and we denote

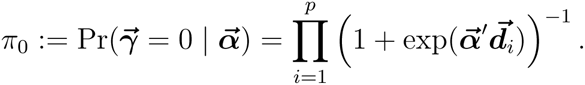

We use 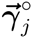 to denote the single-SNP association model where the j-th SNP is the assumed QTN. Clearly,

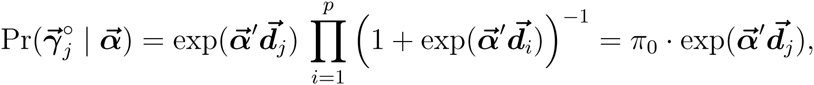

and

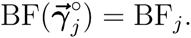

We recall that BF_*j*_ denotes the Bayes factor based on the single-SNP analysis of SNP *j*. The computation of BF_*j*_ has been detailed by many authors.^17,31,32^ It typically requires only summary-level statistics, e.g., the estimated genetic effect of the target SNP and its standard error,^31,32^ and it is computationally trivial.

Finally, we note that given the restrained model space, the PIP of SNP *j*, 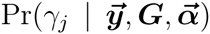, coincides with 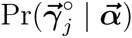. Given all of the above, it follows from simple algebra that

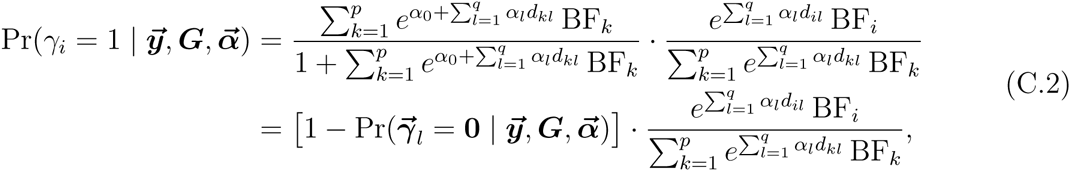

where the first term assesses the probability that the *p*-SNP locus contains a QTL and the second term is the conditional probability that the *i*-th SNP is the sole QTL. The expression (C.2) bears great similarity to the previously proposed Bayesian approaches,^9,11,23^ which also impose the “single QTL per locus” assumption. However, all the aforementioned approaches formulate it as a prior assumption, which results in a very different parametrization. More specifically, they use a locus-level quantity, *π*_0_, to denote the probability that a locus does not contain a QTL. Conditioning on the case that the locus does contain a QTL, the prior for SNP *i* being the causal SNP is assigned

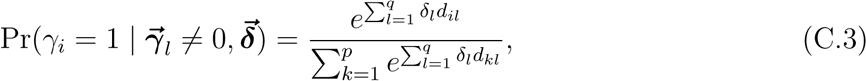

where the parameter 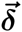 is similar to our enrichment parameter. As a result, this parametrization yields a similar expression for the PIP of SNP i,

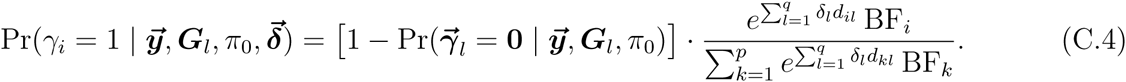

Despite the algebraic similarity, the parameters (*π*_0_ and 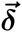) in (C.4) cannot be directly inter-preted as 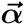 in our logistic priors, partly due to the conditional nature of the prior specification (C.3). Furthermore, in enrichment analysis, the M-step of the EM algorithm becomes much more involved for optimizing the objective function jointly with respect to (*π*_0_, 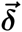). In comparison, we have shown that under the parametrization of DAP-1, the maximization in the M-step is equivalent to fitting a logistic regression model for which the solutions are well known.

## Appendix D Factorization of the posterior probability by LD blocks

For integrative association analysis for loci spanning very large genomic regions, especially in GWAS settings, we recommend an additional approximate factorization, 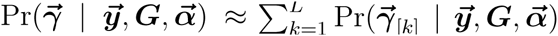, before applying the DAP to each genomic region independently. We provide the necessary mathematical justification for this factorization.

It is sufficient to show that

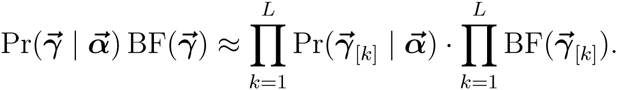

Recall that 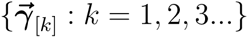 are non-overlapping segments of the vector 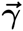. Because the prior probabilities are assumed to be independent across SNPs, it follows trivially that 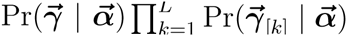.

To show that

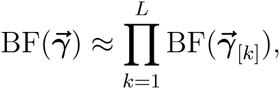

we note the previous result on the Bayes factors,^18^

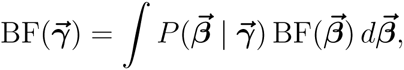

where the probability 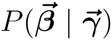 defines the prior effect size given association status 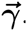. Furthermore, note the independent relationship of the prior effect sizes across SNPs,

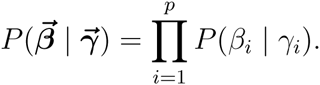

If *γ_i_* = 1, *β_i_* is assigned a normal prior, whereas if *γ_i_* = 0, *β_i_* = 0 with probability 1 (or is represented by a degenerated normal distribution, *β_i_* ~ N(0, 0)). Equivalently, we write

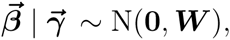

where ***W*** is a diagonal prior variance-covariance matrix, and for 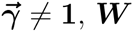 is singular.

Without loss of generality, we assume that both the phenotype vector 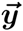 and the genotype vectors 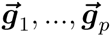 are centered, i.e., the intercept term in the association model is exactly 0. Furthermore, we also assume that the residual error variance parameter *τ* is known. It then follows from the result of Wen^18^ that

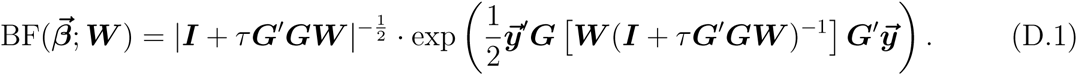

This expression provides the theoretical basis for the factorization. In particular, the *p* × *p* sample covariance matrix 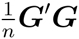 is a well-known estimate of Var(***G***). In other words, ***G′G*** can be viewed as a noisy observation of *n*Var(***G***). Using population genetic theory, Wen and Stephens^24^ show that Var(***G***) is extremely banded. Based on this result, Berisa and Pickrell^25^ recently provided an algorithm to segment the genome into *L* non-overlapping loci utilizing the population parameter of the recombination rate, i.e.,

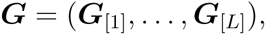

and we approximate ***G'G*** by a block diagonal matrix

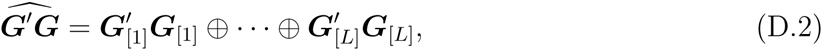

where “⊕” denotes the direct sum of the matrices. It is important to note that (D.2) should be viewed as a de-noised version of ***G'G*** with non-zero entries outside the LD blocks shrunk to exactly 0. By plugging (D.2) into (D.1), it follows that

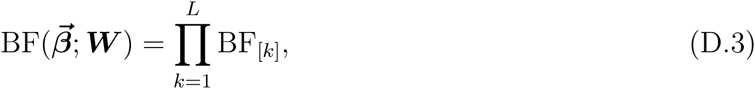

where

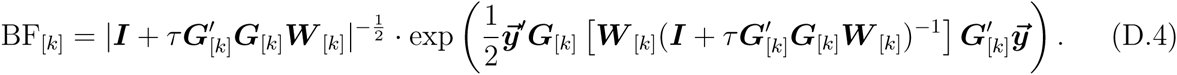

In particular, (***W***_[1]_,…, ***W***_[[*L*]_) is a decomposition of the diagonal matrix ***W*** compatible with the decomposition of ***G***.

Finally, we integrate out the residual error variance parameter *τ* for each BF_[*k*]_ by applying the Laplace approximation.^18^ This step results in plugging in a point estimate of *τ* (e.g., based on 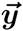 and ***G***_[*k*]_ for each block *k*) into the expression (D.4). Taken together, we have shown that

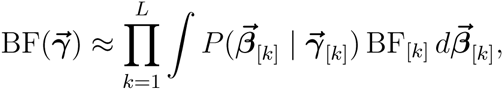

and consequently,

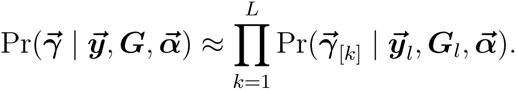

## Appendix E Average Accuracy of PIP Estimates using DAP-1

In this section, we provide some mathematical arguments to justify that DAP-1 (or adaptive DAP with less stringent threshold values) algorithm can provide *on average* accurate estimate. Specifically, we write the expression for the exact calculation of the PIP for SNP *k* at locus *l* as

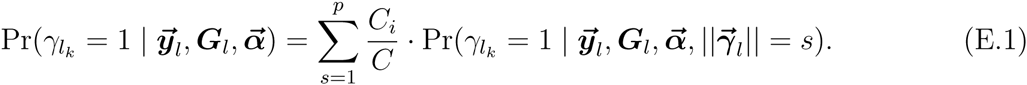

In the case of DAP-1, we essentially use the following expression to approximate the PIP,

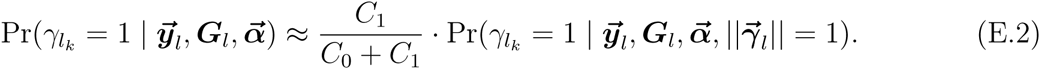

Note that in genetic association analysis, the vast majority of SNPs have overall PIPs → 0 within any given locus; hence, it must be the case that for such a SNP *k*,

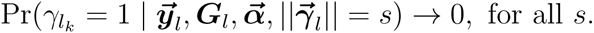

Therefore, even *C*_1_ + * *C*_0_ approximates *C* poorly, and (E.2) still provides an adequately accurate PIP estimation for the majority of SNPs that are not QTNs. The same argument can also be applied to candidate QTNs with very strong evidence for associations, especially when the “primary” association signals have strengths of associations that are orders of magnitude higher than the remaining candidate SNPs within a locus (e.g., 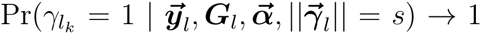 for all *s*). Therefore, the only SNPs whose PIPs are poorly approximated by DAP-1 are those secondary QTL signals (if there are any), but in most practical cases, it can be assured that such SNPs are small in number.

